# RaMALDI: enabling simultaneous Raman and MALDI imaging of the same tissue section

**DOI:** 10.1101/2023.05.07.539107

**Authors:** Ethan Yang, Jeong Hee Kim, Caitlin M. Tressler, Xinyi Elaine Shen, Dalton R. Brown, Cole C. Johnson, Ishan Barman, Kristine Glunde

## Abstract

Multimodal tissue imaging techniques that integrate two complementary modalities are powerful discovery tools for unraveling biological processes and identifying biomarkers of disease. Combining Raman spectroscopic imaging (RSI) and matrix-assisted laser-desorption/ionization (MALDI) mass spectrometry imaging (MSI) to obtain fused images with the advantages of both modalities has the potential of providing spatially resolved, sensitive, and specific biomolecular information, but has so far involved two separate, consecutive tissue sections for RSI and MALDI MSI, resulting in images from two separate entities with inherent disparities. We have developed RaMALDI, a streamlined, integrated, multimodal imaging workflow of RSI and MALDI MSI, performed on a single tissue section with one sample preparation protocol. We show that RaMALDI imaging of various tissues effectively integrates molecular information acquired from both RSI and MALDI MSI of the same sample.

**Table of Contents:** 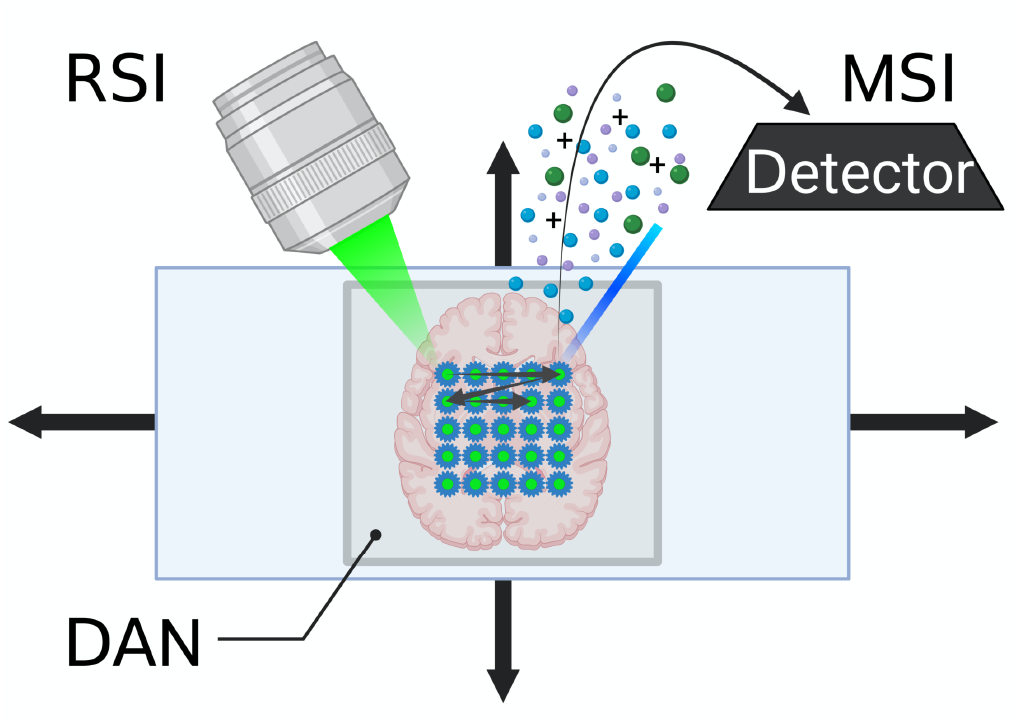

We demonstrate for the first time RaMALDI imaging, a streamlined, integrated multimodal imaging workflow of Raman spectroscopy imaging (RSI) and matrix-assisted laser desorption/ionization mass spectrometry imaging (MALDI MSI), which is performed on a single sample and uses one sample preparation protocol. RaMALDI imaging of various tissues effectively integrates molecular information acquired from both RSI and MALDI MSI of the same sample.

## Introduction

Development of microscopic imaging technologies that probe the anatomical, functional, and molecular features of biological tissues has significantly contributed to biomedical research and clinical applications. However, single imaging modalities are inherently limited in application, whether it be by scope, scale, or sensitivity of the imaging technology, or the preparation and analysis requirements before and after the imaging procedure. Combining imaging techniques that complement one another extends their applications, allowing the investigation, assessment, and monitoring of increasingly diverse and complex biological systems^[1]^. Multimodal imaging that combines the strengths of different modalities has emerged for a more comprehensive characterization of various diseases including cancers and their microenvironment^[2]^. Microscopic tissue images collected from different modalities have the potential to amplify and substantiate information by complementing each other. The use of multiple independent modalities helps to overcome the limitations inherent to individual sensors and reduce noise sources specific to a particular technology. However, while advanced multimodality imaging applications are currently expanding the scope of biomedical research, significant technical challenges still exist for combining certain technologies, including the integration of Raman spectroscopic imaging (RSI) and matrix-assisted laser-desorption/ionization (MALDI) mass spectrometry imaging (MSI).

Raman spectroscopy is widely used to identify biomolecular information in samples with chemical specificity^[3]^. Its label-free and non-destructive detection extends its applications to various biological samples including live cells, tissues, organs, and organisms^[4-6]^. Raman spectroscopy detects vibrational molecular information of a sample, allowing for quantitative analysis of lipids, nucleic acids, and proteins among other molecules^[6-10]^. RSI has the ability to resolve subcellular structures at submicron scales up to the diffraction limit of incident light, increasing its utility for studies that seek to understand the spatial distribution of these biomolecules in tissue samples^[11]^. However, RSI quality can be affected by fluorescence background from biomolecules or impurities of the sample^[12]^. It also suffers from relatively poor sensitivity due to weak scattering efficiency, making imaging time rather long^[13]^. For example, mapping 800×800 μm^2^ with a spatial resolution of 100 μm at an acquisition time of 0.5 sec will require about 2.3 minutes; however, if we desire an improved spatial resolution of 10 μm, then the acquisition time for RSI increases to almost 3.7 hours. This prolonged acquisition time could lead to sample degradation during the time of imaging.

MSI, on the other hand, provides a much greater level of sensitivity than RSI. Soft ionization techniques, such as MALDI MSI, are able to quantitatively detect and resolve tens to hundreds of *m/z* peaks in one experiment, each corresponding to a unique molecule^[14-15]^. Recent advances in laser technology and sample preparation techniques have enabled faster detection (mapping 800×800 μm^2^ at 100 μm spatial resolution requires about 8 seconds and at 10 μm spatial resolution about 7 minutes) of numerous classes of biologically relevant molecules at low micron resolution, ranging from small metabolites^[16]^ and drugs^[17]^ to lipids^[18-19]^, gangliosides^[20-21]^ and glycans^[22-23]^, making MALDI imaging an important tool in preclinical studies^[24]^. While submicron MALDI imaging has been reported in the literature^[25]^, the current state-of-the-art in MALDI matrix deposition and MSI instrumentation does not allow for spatial resolutions below 5 micron^[26-27]^. Combining RSI and MALDI MSI in a multimodal imaging approach would considerably enhance the molecular specificity and spatial resolution of multiplexed, microscopic tissue imaging.

To achieve this goal, there have been a few studies on combining RSI and MALDI MSI to extract a condensed image with the advantages of both modalities. However, all previous studies involved two separate, consecutive tissue sections – one for RSI and the other one for MALDI MSI^[28-29]^. Moreover, some studies were limited to spectral comparisons^[28-32]^. For producing correlated images, post-processing approaches were applied which required image conversion^[33]^ and extensive image registration^[34-35]^. Although these studies reported some correspondence between RSI and MALDI MSI images collected from two consecutive sections, they were essentially limited by not being obtained from the same cells and tissues architectures across both methods, as each tissue section is separated from its adjacent sections by approximately 10 μm determined by the thickness of tissue sectioning^[36]^. RSI and MALDI MSI of the same sample using a single sample preparation method remains unexplored because of perceived compatibility issues between substrates and differences in sample preparation protocols. One recent study showed an integrated workflow with a surface-enhanced mode of MSI and RSI, which, however, requires a specialized nanostructured substrate for measurement^[37]^.

Here, we report the development of RaMALDI, a novel workflow that seamlessly integrates RSI and MALDI MSI of a single sample. Prior to our work, it was unclear whether this integration would be possible because of vastly different sample requirements for each technique. However, we hypothesized that a suitable substrate might facilitate the integration of the two techniques. After trials involving many matrix and sample preparation candidates, we identified a protocol that uniquely preserved both spectral and spatial qualities of integrated RSI and MALDI MSI. We applied the MALDI matrix 1,5-diaminonaphthalene (DAN) over a sample prepared on an indium-tin oxide (ITO) slide, a substrate typically disregarded in Raman spectroscopy due to spectral interference concerns. This finding led to the establishment of RaMALDI, facilitating multimodal imaging of RSI and MALDI MSI with considerably simplified sample preparation and minimal post-processing. We have demonstrated our approach across progressively complex samples, ranging from a nonbiological sample to liver homogenate, kidney, and brain tissue specimens. The resulting highly multiplexed multimodal images from RaMALDI imaging will allow us to better understand biological processes in a morphological and structural tissue context.

## Results and Discussion

We developed RaMALDI, an integrated workflow of RSI and MALDI MSI as shown in **Fig. 1**. Our approach measures both RSI and MALDI MSI of a single sample. We tested and validated our new approach by using non-biological and several biological samples including liver, kidney, and brain as shown in the following.

**Figure 1.**
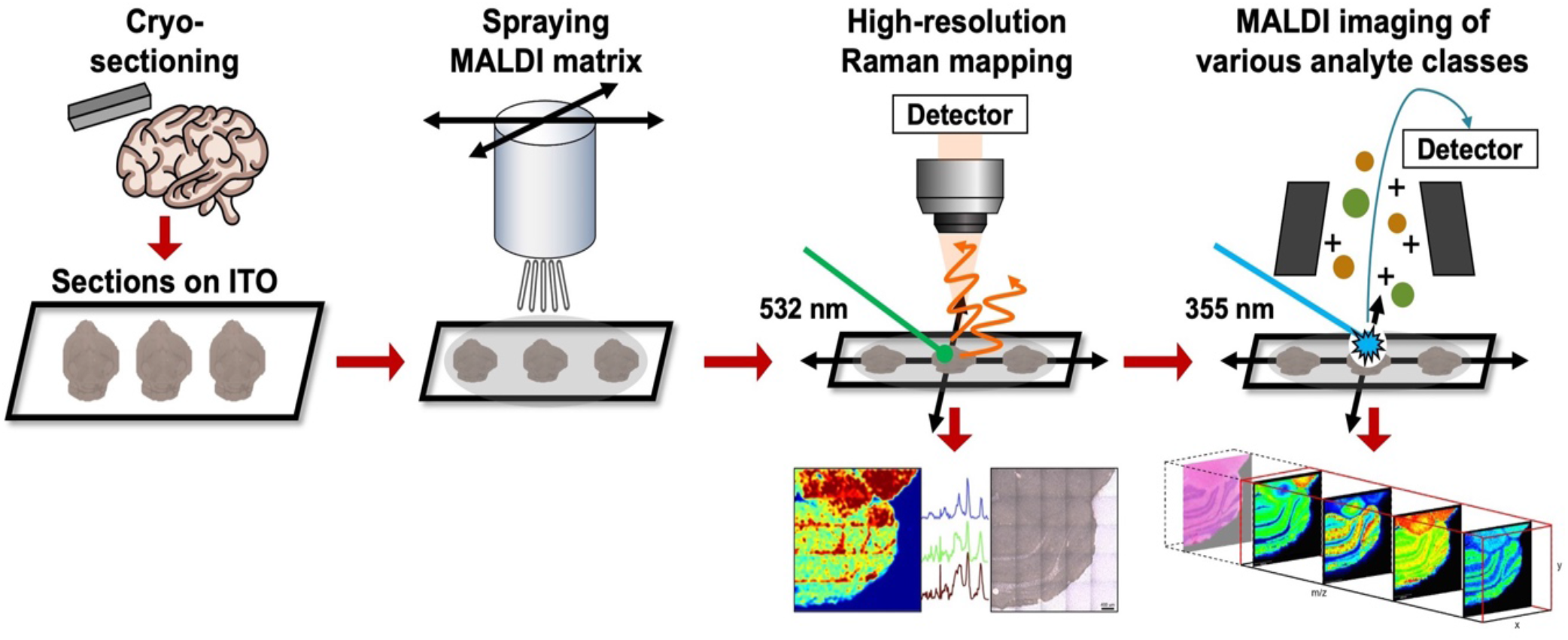
Experimental workflow for RaMALDI imaging. A fresh-frozen tissue section is cryo-sectioned and thaw-mounted onto a conductive ITO microscopy slide. This is followed by MALDI matrix application by spraying DAN onto the tissue section. DAN was identified by an extensive optimization process to have the least influence on spectral quality of RSI and MALDI MSI measurements. The prepared tissue section is first utilized for RSI and then the same sample is subjected to MALDI MSI measurement. This workflow enables the seamless production of two separate images of a single tissue section without compromising the spectral or spatial quality of the hyperspectral datasets.

### RaMALDI Optimization

Optimization was necessary to run both RSI and MALDI MSI measurements on the same slide. Samples are usually prepared on quartz slides for RSI to minimize background (fluorescence and Raman) signal, while MSI frequently uses conductive slides, typically ITO slides, with various types of matrices sprayed on top of the tissue section. We tested and optimized the use of ITO slides with different MALDI matrices for RSI using drawings of *Sharpie****®*** permanent markers, which have high signal-to-noise ratio (SNR) for both RSI and MALDI MSI. We tested 9 different colors of *Sharpie****®*** – pink, brown, red, purple, orange, blue, green, black, and yellow – marked on an ITO slide. Among them, pink presented the clearest signal for both RSI and MSI (**Table S1**), thus we conducted optimization of the workflow with a pink *Sharpie****®*** marker.

RaMALDI was optimized with two goals: (1) enabling both RSI and MSI measurements on a single sample; and (2) crucially, achieving minimal-to-no influence of RSI measurements on the quality of MSI measurements, which has presented a key challenge in employing both modalities on the same sample. For this purpose, we tested the potential suitability of various MALDI matrices based on the imaging performance. While spontaneous Raman spectroscopy typically does not require any foreign agents for measurement, matrix deposition is a necessary step for MALDI MSI to ionize molecules. We measured Raman spectra of pink *Sharpie****®*** marks on an ITO slide typically used for MALDI MSI, sprayed with the six most common MALDI matrices 1,5-diaminonapthalene (DAN), alpha-cyano-4-hydroxycinnamic (CHCA), 2,5-dihydroxybenzoic acid (DHB), 9-aminoacridine (9AA), norharmane (nH), and sinapinic acid (SA) (**Fig. S1**). Notably, Raman spectra acquired from DAN-sprayed pink *Sharpie****®*** marks on ITO slide showed almost no difference when compared to Raman spectra acquired from pink *Sharpie****®*** marks on ITO slide without matrix, or compared to *Sharpie****®*** marks on quartz slide, the latter of which is a standard setup for Raman measurements (**Fig. S1C**). Next, we examined if the matrix density of DAN deposition had any effect on the Raman spectra. Raman peaks of the pink *Sharpie****®*** marker were preserved regardless of DAN matrix density (**Fig. S2**). Based on these findings and since the optimal DAN matrix density for MALDI MSI of biological samples is 0.0016 mg/mm^2^, we selected this same DAN density for subsequent RaMALDI measurements. All RaMALDI experiments were conducted with the optimized setup of sample prepared on an ITO slide, followed by spraying DAN at 0.0016 mg/mm^2^ density over the sample, followed by RSI, and then MALDI MSI, as shown in the workflow in **Fig. 1**.

### Non-biological Sample: Sharpie® Permanent Marker

We tested the RaMALDI imaging workflow with simple, nonbiological models of various *Sharpie****®*** permanent marker drawings in the shape of a “**J**”. RaMALDI was acquired from 9 different *Sharpie****®*** colors – pink, brown, red, purple, orange, blue, green, black, and yellow. While all colors clearly showed “**J**” markings with distinctive RSI and MALDI MSI peaks, pink, brown, and red *Sharpie****®*** markers showed the highest SNR for both the RSI and MALDI MSI portions of RaMALDI imaging (**Fig. 2**), which also agreed with the results of the RaMALDI optimization (**Table S1**). RaMALDI RSI images were mapped to the spatial distribution of baseline corrected, normalized Raman intensity based on the identified peaks unique to each color, which were 1650, 1406, and 621 cm^-1^ for pink, brown, and red *Sharpie****®*** markers, respectively. The observed Raman peak at 1650 cm^-1^ in the pink marker was attributed to C=C stretching vibration of the xanthene ring system of Rhodamine B^[38]^ and Rhodamine 6G^[39]^ and the quinoid ring system of quinacridone^[40]^, which are commonly used for pink pigments. The characteristic peak at 1406 cm^-1^ in brown likely resulted from carbon-based molecules, including quinacridone^[40]^, azo compounds^[41]^, and isoindolinone^[42]^. In red, the peak at 621 cm^-1^ was attributed to azo compounds^[43]^, naphthol^[43-44]^, and phthalocyanine^[44]^. Similarly, RaMALDI MSI data were displayed at the corresponding m/z signals of m/z^+^ 443.4, m/z^-^ 649.1, and m/z^+^ 415.3, which were unique for pink, brown, and red *Sharpie****®*** markers, respectively. The observed peak at m/z^+^ 443.4 in pink marker and m/z^+^ 415.3 in red marker resulted from Rhodamine 6G and its derivatives^[45-46]^. The distinct peak at m/z^-^ 649.1 was likely caused by red dyes, such as direct red 28 and acid red 87 from the Schweppe collection^[46]^. Through peaks specific to each marker color as measured by RaMALDI RSI and MSI, we were able to identify Rhodamine B and Rhodamine 6G in pink marker, azo compounds in brown marker, and naphthol and Rhodamine 6G in red marker, which is consistent with previous studies using Raman and mass spectrometry-based identification of color pigments^[47]^. Along with the spectral domain, spatial images of both RaMALDI RSI and MSI clearly show the “**J**” markings with a uniform signal distribution and clear distinction from the background for each color-specific Raman and *m/z* peak. The corresponding spectra of all measured *Sharpie****®*** marker colors for both RSI and MALDI MSI are shown in **Fig. S3-S5**.

**Figure 2.**
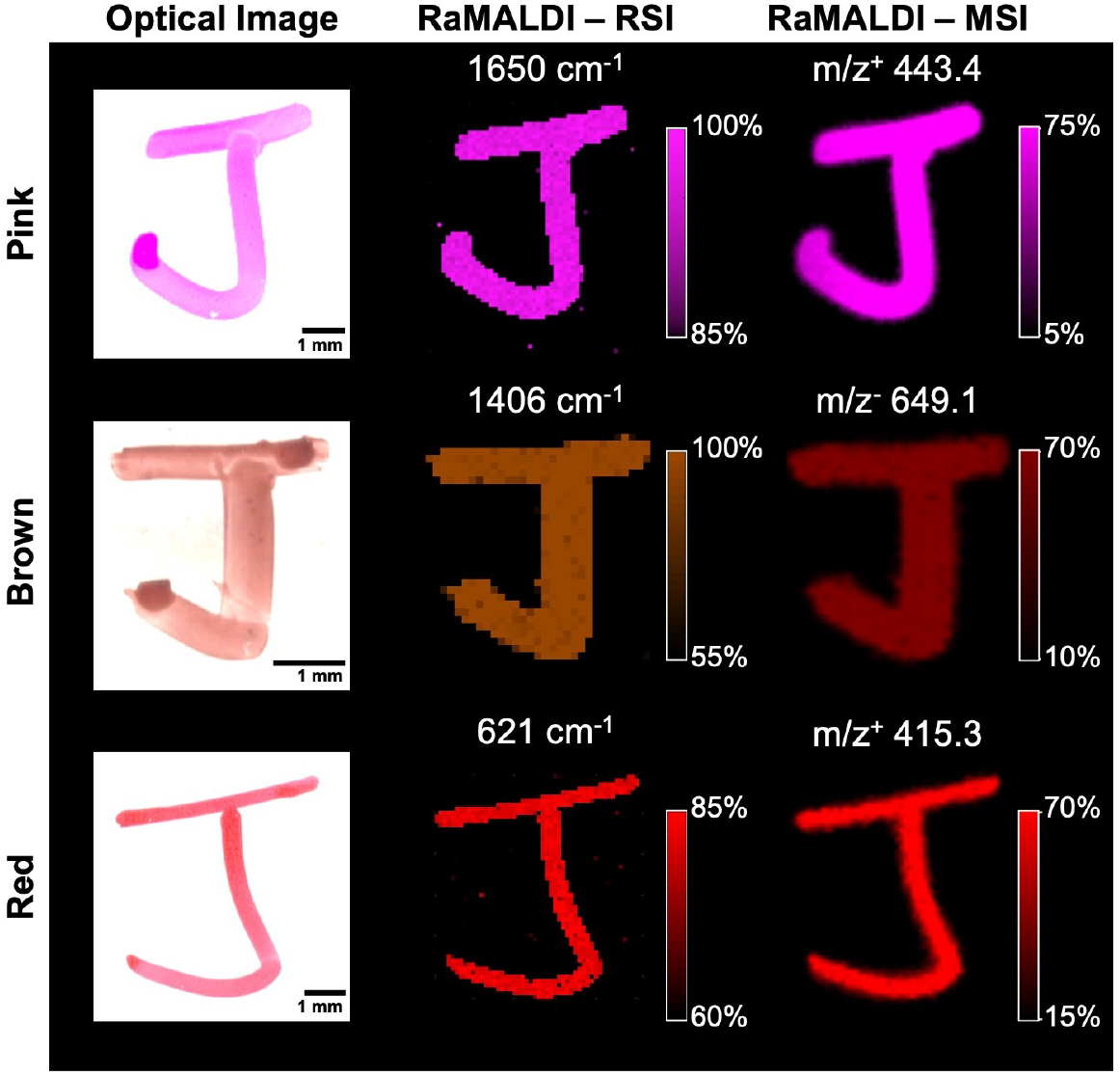
RaMALDI imaging of Sharpie® markers. Pink, Brown, and Red (from top to bottom) markings on an ITO slide underneath 0.0016 mg/mm^2^ DAN matrix were measured using RaMALDI imaging. Both RSI and MSI of RaMALDI imaging were individually processed and images reconstructed at unique Raman and *m/z* peaks, respectively, for each Sharpie® marker color. Identified Raman and *m/z* peaks: 1650 cm^-1^ and m/z^+^ 443.4 for pink, 1406 cm^-1^ and m/z^-^ 649.1 for brown, and 621 cm^-1^ and m/z^+^ 415.3 for red, were selected as the most prominent peaks compared to background signal. Raman peaks at 1650 cm^-1^, 1406 cm^-1^, and 621 cm^-1^ mainly arise from Rhodamine B and 6G, azo compounds, and naphthol and Rhodamine 6G for pink, brown, and red pigments, respectively. MSI peaks at m/z^+^ 443.4 and m/z^+^ 415.3 result from Rhodamine 6G and its derivatives. The MSI peak at m/z^-^ 649.1 arises from red dyes, such as direct red 28, frequently used in brown-colored markers. Scale bar = 1 mm.

### Biological Sample: Liver Tissue Homogenate

With the established workflow optimized using nonbiological samples, we tested RaMALDI using cryosections from tissue homogenates of mouse liver tissue. Liver tissue homogenate was selected because it provides characteristic tissue signals for a cross-section without characteristic morphological structures, as recently established in tissue mimetic model development for quantitative drug imaging using MALDI MSI^[48]^. We measured both Raman and MALDI mass spectra using the RaMALDI workflow with DAN matrix sprayed onto a liver tissue homogenate cryosection that was thaw-mounted onto an ITO slide (**Fig. S6A**). Both Raman and mass spectra showed the presence of various lipids in the sample. Using consecutive liver homogenate sections, we compared Raman spectra from the RaMALDI workflow to Raman spectra from standard Raman microspectroscopy, in which the consecutive liver homogenate section was placed on a separate quartz slide without DAN matrix (**Fig. S6B**). Identical lipid peaks were detected in Raman spectra from liver tissue homogenate sections measured in the RaMALDI workflow as compared to those measured with standard Raman spectroscopy, demonstrating the feasibility of the RaMALDI workflow for biological applications without compromising Raman signal integrity and quality. Additionally, we discovered that spraying DAN matrix onto biological samples protects these samples from degradation (**Fig. S6C**). This was evident from observing the characteristic amide-II Raman peak at 1586 cm^-1^ in liver tissue homogenate sections over time. While a decrease in the 1586 cm^-1^ peak intensity was observed for a sample without DAN matrix sprayed onto the sample, the same peak remained at the initial intensity level when measuring a sample covered with DAN matrix for at least 72 hours, which was the duration of our longitudinal monitoring. This indicates that DAN matrix deposition onto biological tissue sections protects from direct exposure to the environment and laser-induced thermal and photodegradation during RSI measurements. Hence, this protective effect of DAN matrix for biological samples is beneficial for long-duration imaging tasks. For tissue imaging, especially at a high spatial resolution, RSI measurement can take several hours depending on the acquisition setup and mapping coverage, making it challenging to guarantee unaltered sample quality until MALDI MSI measurement is performed. This is a significant drawback in all previously reported studies for combining RSI and MALDI MSI on the same sample, which were all conducted in the order of RSI, matrix deposition, and MALDI MSI measurement^[34, 49-50]^.

### Biological Sample: Kidney Tissue

Moving into whole tissues, we tested the performance of RaMALDI imaging using tissue sections from fresh-frozen mouse kidneys. **Fig. 3** shows Raman and MALDI MS images of the same kidney tissue section collected using the RaMALDI workflow, along with the corresponding hematoxylin and eosin (H&E) stained image (**Fig. 3A**) from the same section measured following RaMALDI imaging for identification of renal anatomical structures. In the top row of **Fig. 3C**, the RSI data of RaMALDI imaging of kidney tissue sections show the spatial distribution of two resolved chemical species. In particular, these represent the spatial distribution of components 1 and 2 identified using a spectral unmixing technique, termed multivariate curve resolution–alternating least squares (MCR-ALS). Detailed information regarding the generation of spatially resolved chemical images based on MCR-ALS modeling of Raman hyperspectral data is provided in the Supporting Information. **Fig. 3D** shows the resulting chemical spectra from MCR-ALS model-decomposed RaMALDI RSI data, including the Raman peaks that correspond to assignments listed in **Table S2**. For component 1, Raman peaks at 1372, 1437, 1456, and 1575 cm^-1^ were observed, which mostly resulted from nucleic acids^[7]^. Also, the renal capsule was visualized on the chemical Raman map, which resulted from its high content of collagen and elastin in the fibrous extracellular matrix^[51]^. Raman peaks for component 2 included 1366, 1372, 1437, and 1445 cm^-1^, which mainly arose from lipids^[7]^. The identified Raman peaks are consistent with those reported in previous studies of the kidney under various conditions^[52-55]^. While the MCR-ALS analysis of RSI images provides an important starting point for understanding the classes of compositional contributors and their spatial distribution, it does not offer definitive confirmation of the specific molecules in the tissue sample, making the integration with MALDI MSI particularly significant and insightful. The bottom two rows of **Fig. 3C** show both positive and negative ion images of RaMALDI MSI data at selected *m/z* peaks, which were identified as phosphatidylcholine (PC) (36:4), PC (O-38:6), 1-hexadecanoyl-2-(5-oxo-7-carboxy-6E-heptenoyl)-sn-glycero-3-phosphoserine (PKODiA-PS), and phosphatidylserine (PS) (36:2) using on-tissue MS/MS fragmentation for identification of the selected peaks as shown in **Fig. S7-S10. Fig. 3E** shows the corresponding whole tissue mass spectra of RaMALDI MSI data including the m/z signals displayed as MS images in **Fig. 3C**. RaMALDI RSI and MSI images from the same kidney tissue section showed clear spatial correspondence of anatomical structures including renal cortex, medulla, and capsule, which were particularly apparent in the MSI segmentation images (**Fig. 3B**).

**Figure 3.**
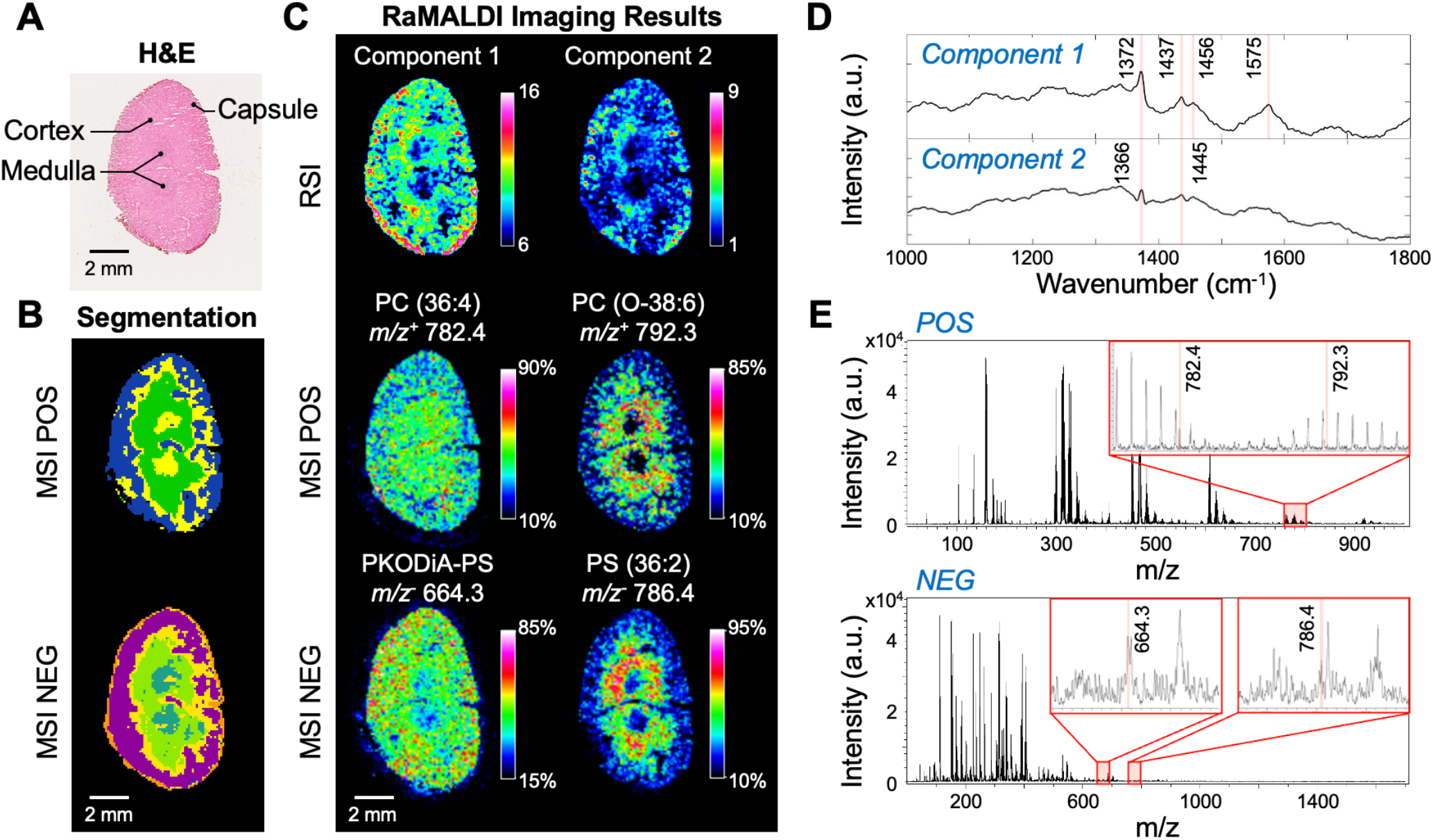
RaMALDI imaging of mouse kidney tissue section. RaMALDI imaging results are shown for RSI, followed by MSI of a representative coronal mouse kidney tissue section on an ITO slide sprayed with DAN matrix and imaged at 100 μm spatial resolution. (**A**) H&E-stained image of the same kidney tissue section was acquired following RaMALDI imaging. Anatomical structures of renal cortex, renal medulla, and capsule are labeled in the image. (**B**) K-means bisecting segmentation results of RaMALDI MSI data from positive (POS) and negative (NEG) MSI of the same section. (**C**) RSI data (top row) and MSI data (middle and bottom rows) of RaMALDI imaging. Two chemical component images (Component 1, Component 2) from the MCR-ALS model are displayed for RSI data. MALDI MS images are shown for two *m/z*’s for positive ion mode identified as specific phosphatidylcholines (PC) and two *m/z*’s for negative ion mode identified as specific phosphatidylserines (PS). Scale bar = 2 mm. (**D**) MCR-ALS resolved RaMALDI-RSI spectra of component 1 (nucleic acid-rich) and component 2 (lipid-rich), corresponding to (C) RaMALDI-RSI. Pure chemical spectra were decomposed from preprocessed RaMALDI-RSI spectra in the biological fingerprint region (1000-1800 cm^-1^) based on MCR-ALS model corresponding to (C). (**E**) RaMALDI-MSI spectra of positive ion (top) and negative ion (bottom) modes corresponding to (C). M/z signals shown as images in (C) are highlighted in red in the expanded spectral regions.

### Biological Sample: Brain Tissue

We further tested high spatial resolution RaMALDI imaging of fresh-frozen mouse brain sections. Since the cerebellum has a distinct anatomical and morphological structure with multiple molecular layers, we evaluated RaMALDI imaging for delineating this intricate structure. Brain tissue sections were imaged using the RaMALDI imaging workflow at spatial resolutions of 50 μm (**Fig. 4**) and 5 μm (**Fig. 5**) to evaluate the correspondence between RSI and MALDI MSI images regarding the fine histological features of the cerebellum. H&E-stained histological images were obtained from the same brain tissue section following RaMALDI imaging and are shown in **Fig. 4A** and **5A** to identify distinct anatomical structures in the cerebellum. The gray and white matter were clearly distinguished in both RSI and MSI images of RaMALDI imaging as shown in **Fig. 4C**. The gray matter, which is mainly composed of myelinated axons of granule cells and dendrites of Purkinje cells, showed Raman peaks from both lipid and protein, identified in **Fig. 4D** and **5C**. The white matter also displayed lipid and protein signatures from myelinated neuronal axons, and peaks associated with nucleic acids from cerebellar nuclei. These vibrational peaks were previously reported in studies of brain tissues with Raman microspectroscopy ^[56-59]^. The bottom two rows of **Fig. 4C** display positive and negative ion modes of MALDI MSI images at selected, identified *m/z* peaks in **Fig. 4C**. Using on-tissue MS/MS fragmentation analysis of the selected peaks, as shown in **Fig. S11-S14**, we identified these lipids to be PC (38:6), PC (40:4), phosphatidylinositol (PI) (18:0/20:4), and C24:1 sulfatide (cis-tetracosenoyl sulfatide). C24:1 sulfatide was reported in previous brain tissue MALDI MSI studies with a similar spatial distribution^[60]^. Previous studies^[61-62]^ have also reported similar anatomical distributions of the detected PC (38:6) and PC (40:4) in white and gray matter while the exact physiological roles of these two PC species remain to be determined. The localization of the detected PI (18:0/20:4) to gray matter suggests a possibly greater role in inter-neuronal signaling than that of other PI species^[62]^. Lastly, sulfatides including C24:1, are mostly found in white matter because they are a major component of the myelin that surrounds nerve fibers^[62]^.

**Figure 4.**
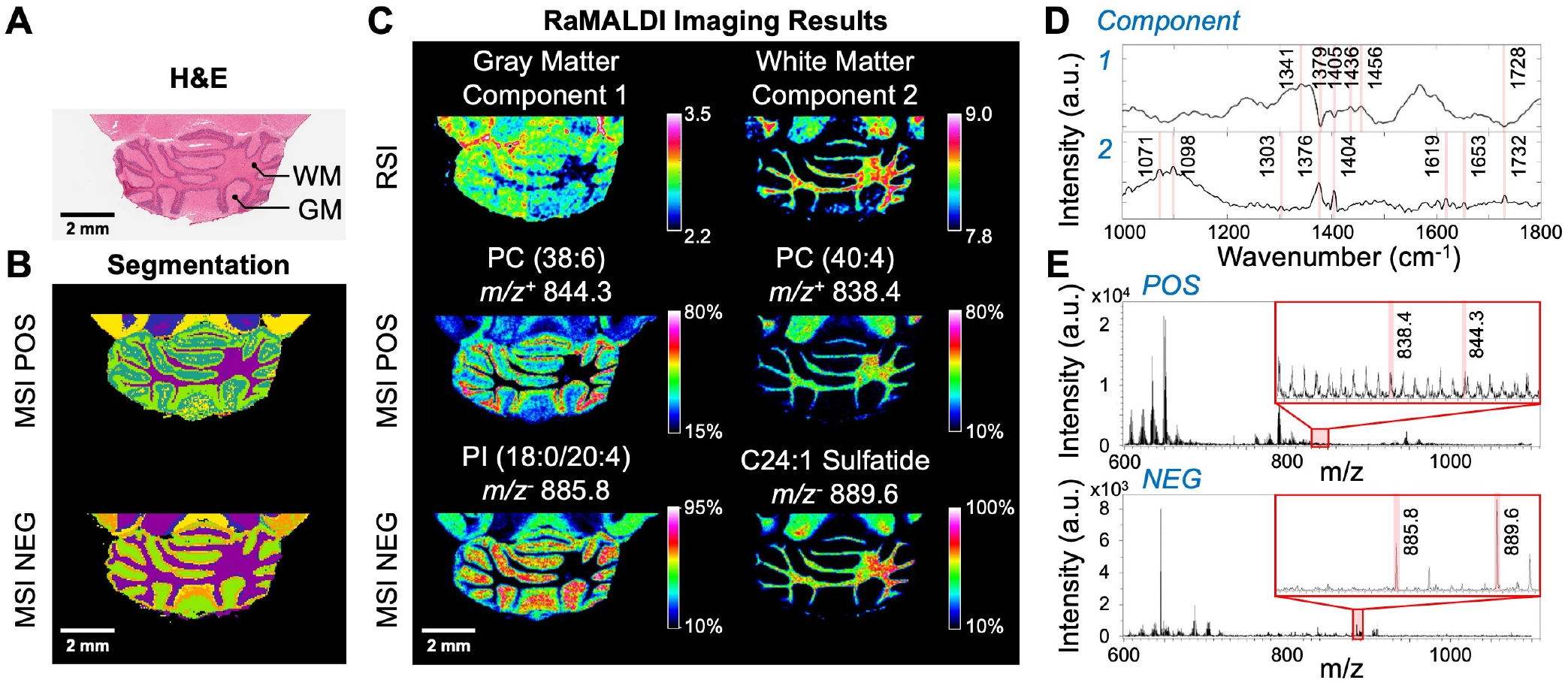
RaMALDI imaging of brain tissue section at 50 μm lateral spatial resolution. RaMALDI imaging results are shown for RSI, followed by MSI of a horizontal brain tissue section on an ITO slide sprayed with DAN matrix and imaged at 50 μm spatial resolution in the region of the cerebellum. (**A**) H&E image of brain tissue section showing gray matter (GM) and white matter (WM) as labeled in the image. (**B**) K-means bisecting segmentation results of RaMALDI MSI data from positive (POS) and negative (NEG) MSI of the same section. (**C**) RSI data (top row) and MSI data (middle and bottom row) of RaMALDI imaging. Two chemical component images (Component 1, Component 2) from the MCR-ALS model are displayed for RSI data. MALDI MS images are shown for two *m/z*’s for positive ion mode identified as specific phosphatidylcholines (PC) and two *m/z*’s for negative ion mode identified as a specific phosphatidylinositol (PI) and a specific sulfatide. Scale bar = 2 mm. (**D**) MCR-ALS resolved RaMALDI-RSI spectra of component 1 (gray matter) and component 2 (white matter). Pure chemical spectra were decomposed from preprocessed RaMALDI-RSI spectra in the biological fingerprint region (1000-1800 cm-1) based on the MCR-ALS model corresponding to (C). (**E**) RaMALDI-MSI spectra of positive ion (top) and negative ion (bottom) modes corresponding to (C). M/z signals displayed as images in (C) are highlighted in red in the expanded spectral regions.

**Figure 5.**
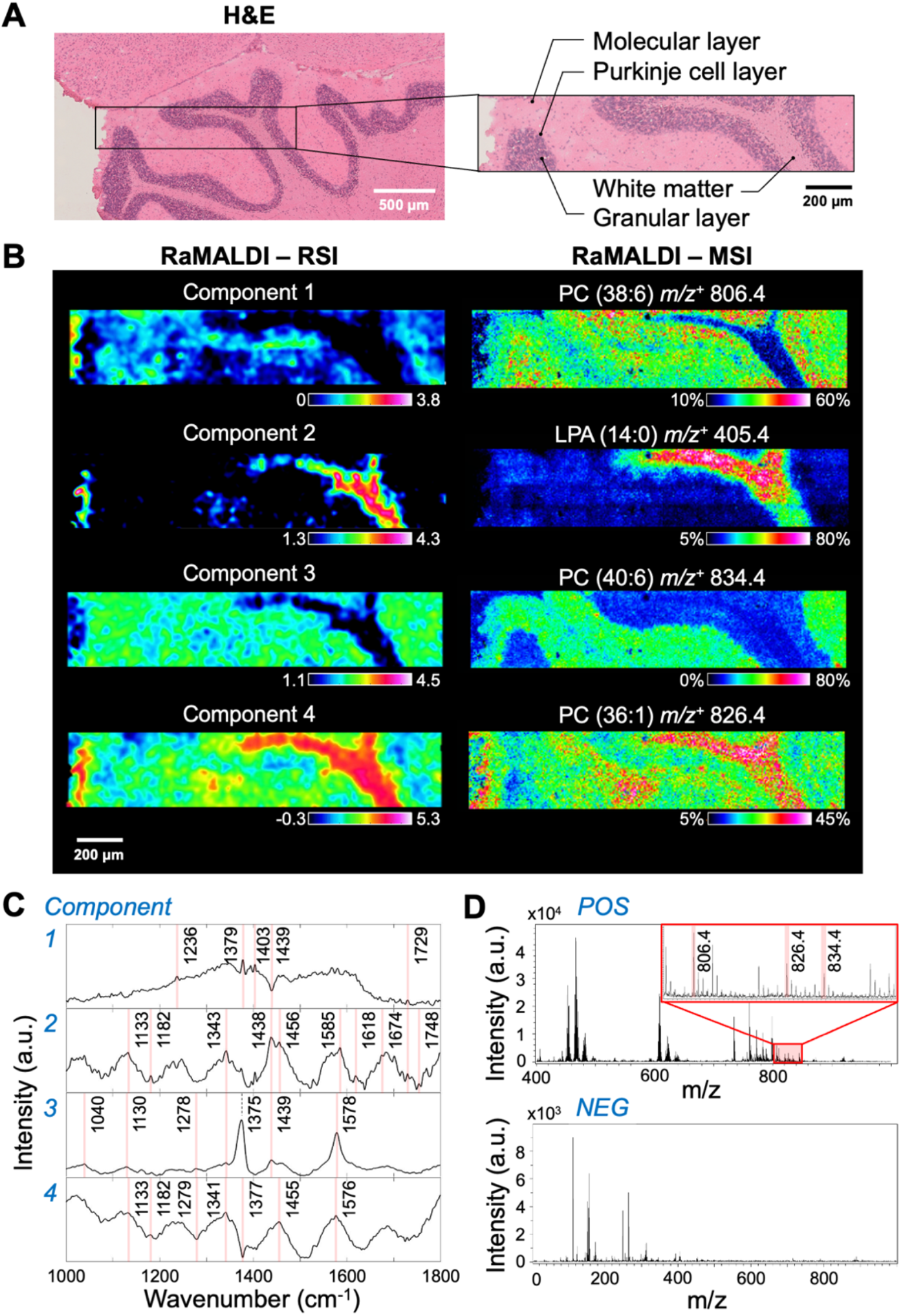
RaMALDI imaging of brain tissue section at 5 μm lateral spatial resolution. RaMALDI imaging results are shown for RSI, followed by MSI of a horizontal brain tissue section on an ITO slide sprayed with DAN matrix and imaged at 5 μm spatial resolution in the region of the cerebellum. (**A**) H&E images of brain tissue section showing the three layers of molecular, Purkinje, and granular layer in gray matter and white matter as labeled in the image. Scale bar = 500 μm (left) and 200 μm (right). (**B**) RSI data (left) and MSI data (right) of RaMALDI imaging. Four chemical component images (Component 1, Component 2, Component 3, Component 4) from the MCR-ALS model are displayed for RSI data. MALDI MS images are shown for four *m/z*’s for positive ion mode identified as three specific phosphatidylcholines (PC) and a specific lysophosphatidic acid (LPA). Scale bar = 200 μm. (**C**) MCR-ALS resolved RaMALDI-RSI spectra of component 1 (gray matter), component 2 (white matter, cholesterol- and lipid-rich), component 3 (gray matter), and component 4 (white matter, lipid-rich). Pure chemical spectra were decomposed from preprocessed RaMALDI-RSI spectra in the biological fingerprint region (1000-1800 cm^-1^) based on the MCR-ALS model corresponding to (B). (**D**) RaMALDI-MSI spectra of positive ion (top) and negative ion (bottom) modes corresponding to (B). M/z signals shown as images in (B) are highlighted in red in the expanded spectral regions.

In the RaMALDI images of brain tissue sections acquired at 5 μm spatial resolution, the molecular distributions, particularly those of lipids, proteins, and nucleic acids, clearly show the molecular layers of the cerebellum. MCR-ALS resolved RSI images show relatively intense signals from components 1 and 3 in gray matter and from components 2 and 4 in white matter as shown in **Fig. 5B** with their respective spectra displayed in **Fig. 5C**. RSI spectra of all components in **Fig. 4D** and **Fig. 5C** show mixed features of nucleic acids, lipids, and proteins. As white matter has a relatively high lipid content, peaks at 1133, 1438, and 1674 cm^-1^, associated with C-C stretch, CH_2_ deformation, and C=C stretch vibrations of lipids were observed in components 2 and 4. Component 2 displayed cholesterol peaks at 1133, 1438, and 1674 cm^-1^, which was previously shown to be one of the most abundant lipids in the brain^[56]^. Also, both components 3 and 4 featured overall signal from the outer molecular layer surrounding the cerebellum. Peaks contributed by protein content in brain tissues were observed in amide I, II, and III bands ranging between 1600-1700, 1480-1575, and 1200-1300 cm^-1^, respectively. All observed RSI peaks of brain tissues and their molecular assignments are summarized in **Table S3**. The right column of **Fig. 5B** shows positive ion mode MSI images at selected, identified *m/z* peaks, whose spectral domain is shown in **Fig. 5E**. Using on-tissue MS/MS fragmentation analysis of these selected peaks, we identified these lipids to be PC (38:6), lysophosphatidic acid (LPA) (14:0), PC (40:6), and PC (36:1) as shown in **Fig. S15-S18**. Previous studies have reported similar anatomical distributions of these three PCs in gray and white matter^[61-62]^, with PC (38:6) localizing to molecular, Purkinje cell, and granular layers, PC (40:6) enriched in the molecular layer, and PC (36:1) detected predominantly in white matter. The detected LPA (14:0) is primarily localized to white matter, suggesting its possible roles in neural activities specific to this region of the cerebellum.

In summary, RaMALDI imaging is a new, streamlined workflow for acquiring highly multiplexed molecular tissue images by performing RSI and MALDI MSI on the same tissue sample. This approach allows for multimodality imaging to characterize molecular features within their respective morphological-anatomical tissue structures. The presented workflow is a crucial step towards rapid, simple multimodal tissue imaging that can be applied in studies aimed at biomedical discovery of clinically relevant biomarkers. While our rationally designed proof-of-principle study has focused on the MALDI matrix DAN, which was based on our discoveries of the lack of RSI signals from DAN and its protection against tissue degradation, future follow-up studies could further explore other MALDI matrices and additional types of tissue samples in healthy and various pathological states.

## Conclusions

We have developed RaMALDI imaging, an integrated workflow of RSI and MALDI MSI conducted on a single sample with a one-step sample preparation method. RaMALDI imaging is advantageous because it allows for analysis of a single tissue section with both modalities, which reduces the amount of sample needed, and significantly streamlines the multimodal approach, ultimately enabling highly multiplexed molecular imaging discoveries in tissue- and cell-based biomedical research.

We have demonstrated RaMALDI imaging applications of nonbiological and biological samples using *Sharpie****®*** marker drawings, cryosections of mouse liver tissue homogenates, mouse kidneys, and mouse brains. We were able to collect spectral and spatial information of an identical tissue section in a robust and reliable manner, without compromising the quality of either. The RaMALDI imaging approach, which measured a single tissue section by consecutive RSI followed by MSI, generated highly multiplexed molecular maps of mouse brain tissue with a lateral spatial resolution of down to 5 μm for both modalities. The RaMALDI workflow offers a significantly simplified data processing and analysis approach, which facilitates effective interpretation of the two multimodal images. Our workflow does not require extensive post-acquisition image registration between RSI and MSI images for analysis, as RaMALDI images are collected from an identical sample prepared in an identical manner, making them inherently registered. Enabling multimodal imaging of a single tissue section significantly simplifies the sample preparation procedure and avoids potential challenges with morphological discrepancies arising from either intentional or unintentional variations in sample preparation protocols. Such morphological variance between two separate, consecutive tissue sections prepared for two different imaging modalities often prolongs the analysis procedure and hampers comprehensive molecular interpretation of the imaged tissue. Lastly, we observed that matrix application onto tissue sections is advantageous as it protects from degradation, thereby enabling longer imaging times.

By demonstrating that RaMALDI imaging is feasible, we have effectively established the feasibility of simultaneous Raman and MALDI imaging of the same tissue section, which will become possible once instrumentation for Raman spectroscopy and MALDI imaging has been integrated.

## Acknowledgments

The authors would like to acknowledge the Johns Hopkins Applied Imaging Mass Spectrometry (AIMS) Core facility where all MALDI MSI experiments were performed. We would also like to acknowledge the National Institutes of Health grants R01 CA213492, R01 CA213428, R01 CA264901, S10 OD030500, 2-P41-EB015871 and DP2GM128198.

