## Supporting Information for "RaMALDI: enabling simultaneous Raman and MALDI imaging of the same tissue section"

[c] Prof. K. Glunde
Departments of Oncology and Biological Chemistry
The Johns Hopkins University School of Medicine
Baltimore, MD, USA 21287

### These authors contributed equally to this work.

* These authors serve as co-corresponding authors.

Supporting information for this article is available.

**Keywords:** multimodal imaging • Raman spectroscopy • mass spectrometry imaging • MALDI • matrix

#### Experimental Procedures

##### Sample Preparation

**Solvents and chemicals.** Unless otherwise noted, all solvents and trifluoroacetic acid (TFA) were purchased from Sigma Aldrich (St. Louis, MO). 1,5-Diaminonapthalene (DAN), α-cyano-4-hydroxycinnamic acid (CHCA), 2,5-dihydroxybenzoic acid (DHB), norharmane (nH), 9-aminoacridine (9AA) and sinapic acid (SA) MALDI matrices were purchased from Sigma Aldrich (St. Louis, MO). Solvents were all MS grades.

**Sample preparation.** For each RaMALDI imaging experiment, specimens were prepared on indium tin oxide (ITO)-coated glass slides (Delta Technologies, Loveland, CO) and used for both Raman spectroscopy and MALDI MSI measurements. The standard approach for Raman spectroscopy was used for comparison, which was to place specimens on quartz slides to minimize spectral interference from slide background. We utilized both nonbiological ("J" drawings of *Sharpie*® permanent markers) and biological (mouse liver, kidney, and brain tissue) samples for the optimization and testing of the RaMALDI imaging workflow.

***Matrix deposition:*** For matrix deposition, 50 mM solutions of each matrix in 1:1 ACN:H2O + 0.1% TFA were sprayed onto the sample with an HTX M5 Sprayer (HTX Imaging, Chapel Hill, NC) using the following parameters: 4 passes, 100 µL/min flow rate, 1200mm/min nozzle/track velocity, 3 mm track spacing, 30°C nozzle temperature, criss cross pattern, and 0 s dry time. For matrix density optimization, all parameters were kept constant, except for the number of passes, which varied from 0 to 8 passes to achieve 0, 0.0004, 0.0008, 0.0012, 0.0016, and 0.0032 mg/mm$^2$ of matrix density.

***Nonbiological samples:*** For *Sharpie*® drawings, 9 different colors - pink, brown, red, purple, orange, blue, green, black, and yellow- were used. We wrote "J" with a size of approximately 5 mm by 5 mm onto ITO slides, onto which we then sprayed the matrix of choice.

***Biological samples:*** All animal experiments were approved by the Institutional Animal Care and Use Committee (IACUC) of the Johns Hopkins University School of Medicine, which is fully accredited by the American Association for the Accreditation of Laboratory Animal Care (AAALAC). Adult athymic nude nu/nu mice (Taconic Biosciences, Rensselaer, NY) of about three months of age were sacrificed, and their kidneys, livers, and brains were frozen in liquid nitrogen vapors. These frozen organs were stored at -80°C until use.

***Homogenized liver samples:*** Liver homogenates were prepared by using four to five livers, which were ground on liquid nitrogen, and 300 mg of tissue were weighed into Precellys bead homogenizer tubes (VWR, Radnor, Pennsylvania, USA) and homogenized using a Bertin Precellys Evolution Homogenizer (Bertin Corp, Rockville, MD) with the following parameters: 2mL, 6800 RPM, 4x30 seconds cycle with 45 seconds pauses. These samples were centrifuged in an Eppendorf 5424R Centrifuge (Eppendorf, Hamburg, Germany) for 1 min at 15000 rpm and frozen into plugs using a home-built silicone mold in liquid nitrogen vapors prior to storage at -80˚C until cryo-sectioning was performed.

***Cryo-sectioning:*** All biological samples including mouse liver homogenates, kidneys, and brains were cryo-sectioned using a Leica CM1860 UV cryostat (Leica Biosystems, Wetzlar, Germany) at 10 µm thickness and thaw-mounted onto ITO slides, followed by spraying of the matrix of choice as described above. Mouse kidneys were sectioned coronally, and mouse brains horizontally.

***Histology:*** For kidney and brain tissue sections, MALDI matrix was washed off following RaMALDI experiments using ethanol for 24 hours, followed by hematoxylin and eosin (H&E) staining conducted on the same tissue section using a standard protocol with Mayer's hematoxylin solution and aqueous Eosin Y solution from Sigma Aldrich (St. Louis, MO). H&E-stained slides were imaged at 40x magnification using a NanoZoomer S210 Digital slide scanner (Hamamatsu Photonics, Shizuoka, Japan). The H&E-stained tissue sections were visualized and exported from Aperio ImageScope (v 12.4.6, Leica Biosystems, Deer Park, IL).

#### Data Acquisition

**Raman spectroscopy and RSI.** Raman spectra of all samples were measured using an XploRA PLUS confocal Raman microspectroscopy (Horiba Jobin-Yvon, Villeneuve-d'Ascq, France) equipped with a 532 nm diode laser for excitation. Raman scattering resulting from light-sample interactions was collected through an objective lens, dispersed with a diffraction grating density of 1800 gr/mm, and recorded by a thermoelectrically cooled CCD camera that was coupled to the microscope.

Experimental parameters varied for each sample. *Sharpie®* markers were raster-scanned with 1% laser power for 0.5s acquisition time for each data point and collected with a 5x objective lens (**Fig. 2**, **Fig. S1**). Kidney tissue sections used 10% laser power for 1s per acquisition point. Raman mapping of kidney tissue (**Fig. 3**), also collected with a 5x objective lens, covered 9 x 6.4 $mm^2$, with a spatial resolution of 100 µm. For mouse brain tissue sections, the larger map with a coverage of 4.4 x 8 $mm^2$ (**Fig. 4**) was raster-scanned at 50 µm spatial resolution with 10% laser power for 0.1s using a 10x objective lens. The mouse brain tissue section acquired with a smaller field of view (**Fig. 5**) was collected at 5 µm spatial resolution with 1% laser power for 0.5s using a 100x objective lens. The Raman spectrum at each acquisition point was obtained by averaging three repetitions for all samples.

**MALDI MSI.** All MALDI MSI experiments were conducted using a rapifleX MALDI TOF/TOF system (Bruker Daltonics, Billerica, MA) equipped with a 10 kHz smartbeam 3D Nd:YAG laser at 355 nm. flexControl (v 4.0) was used to control the instrument and optimize the acquisition parameters, while flexImaging (v 5.0) was used to set up the MALDI imaging experiments and select the regions of interest. The extraction voltage was set to -20 kV, lens voltage of -11.3 kV, reflectron voltages at -20.86 kV, -1.085 kV and -8.6 kV, with a delay time of 100 ns, while the laser fluence was optimized for each matrix. Prior to data acquisition, height adjustment (target profile generation), laser focus tuning, and external mass calibration with red phosphorus were conducted to achieve a mass error <5 PPM.

For *Sharpie®* drawings (**Fig. 2**), data were acquired in either positive or negative ion reflectron mode at 200 laser shots per pixel with a raster width of 100 µm in *m/z* 200-1000 in beam scan mode. For biological tissues, data were acquired using dual polarity. Kidney tissue sections (**Fig. 3**) were imaged using a 100 µm raster width with dual polarity of negative mode from *m/z* 0-1000 followed by positive ion mode from *m/z* 0-1700, using the protocol in[1] without any offset. To achieve this, 50 laser shots were acquired in both modes with beam scan on. Brain tissue sections (**Fig. 4**) were imaged at 50 µm raster width with dual polarity of negative ion mode from *m/z* 0-1000 followed by positive ion mode from *m/z* 0-1700, using the protocol in[1] without any offset, again with 50 laser shots acquired in both modes with beam scan on. For high spatial resolution MSI (**Fig. 5**), brain tissue sections were imaged using a 5 µm raster width with positive ion mode only from *m/z* 400-1000 on the cerebellum. Regions of interest (ROIs) containing areas previously sampled by RSI and without were acquired for comparison. To achieve this, the scanned image was rescaled in ImageJ to 10160 dpi to obtain 2.5 um pixel size and reduce chances of striping. Further, laser shots were optimized to 50 and data acquired with beam scan off to achieve a laser beam size of ≤ 5 µm. MS/MS was conducted with the LIFT unit of the rapifleX TOF/TOF system with argon as a collision gas with an isolation window of ±1 m/z at 60% laser power boost and 4000 laser shots for fragmentation.

#### Data analysis

**Raman spectroscopy and RSI data processing and analysis.** All Raman data were subjected to pre-processing, including noise reduction, and baseline correction followed by normalization[2]. The collected Raman spectra were first denoised with cosmic ray removal and Savitzky-Golay filters to remove the noise while preserving spectral features[3]. The denoised spectra were corrected with a best-fit polynomial-based fluorescence background removal. The spectra were then min-max normalized to correct potential experimental variations. For analysis, the pre-processed Raman spectra were trimmed to biological fingerprint wavenumber region (1,000–1,800 $cm^{-1}$). Kidney and brain tissue spectra were further analyzed with multivariate curve resolution–alternating least squares (MCR-ALS) algorithm to identify constituent compounds in the tissue samples without a priori information of their composition[4]. The MCR-ALS algorithm processed complex Raman spectra at each acquisition point by decomposing unresolved mixtures into pure individual components with respect to their relative concentration. The unmixing of spectra was based on the number of Raman signatures, estimated by singular value decomposition (SVD), to minimize subjectivity of decomposition procedure. The recovery of pure Raman profiles was achieved as MCR-ALS reached convergence through an iterative optimization under the non-negativity constraint on spectra and relative concentration. All Raman spectral data were processed and analyzed by using MATLAB R2021b (MathWorks, Natick, MA).

**MALDI MSI data processing and analysis.** The MALDI MSI data processing varied based on sample type. For nonbiological samples, MALDI MSI data were directly exported from flexImaging (v 5.0, Bruker Daltonics) with total ion current (TIC) normalization. Average spectra for each *Sharpie*® marker were exported from flexImaging and imported into mMass for comparison[5]. For biological samples, MALDI MSI data were imported into SCiLS Lab (v 2021b, Bruker Daltonics) and segmentation analysis was conducted on TIC normalized data through the software’s segmentation pipeline, which conducts peak picking and peak alignment and smoothing prior to segmentation using the bisecting k-means in combination with correlation distance. Segmentation maps were generated and exported from SCiLS Lab. Individual representative m/z features from each segment were identified from the segmentation analysis and the corresponding results were then visualized in flexImaging.

#### Supplementary Tables

**Table S1. Raman and MALDI signal scoring of Sharpie colors.** The Raman and MALDI MSI signals for the 9 Sharpie® markers were scored based on their overall signal intensity, where ***** represents the highest and * the lowest signal intensity and image quality. An overall score was attributed to each color based on their combined Raman and MALDI MSI ranking, where each star represented 1 point, and ranked based on the highest to lowest score. Based on this ranking, the pink Sharpie® marker represented the most suitable color for optimization given the strong and clear signal from both Raman and MALDI MSI data. The corresponding RSI and MALDI MSI spectra and imaging data for this scoring are shown in Fig. 2, S3, S4, and S5.

| **Color** | **Final Score** | **Raman** | **MALDI** | |
|---|---|---|---|---|
| | | Wavenumber | Pos | Neg |
| Pink | 12 | ***** | **** | *** |
| Brown | 11 | ***** | *** | *** |
| Red | 11 | **** | *** | **** |
| Purple | 11 | *** | **** | **** |
| Orange | 10 | *** | *** | **** |
| Blue | 9 | ** | *** | **** |
| Green | 9 | ** | ** | ***** |
| Black | 8 | *** | *** | ** |
| Yellow | 7 | * | *** | *** |

**Table S2. Assignments for Raman peaks associated with DNA/RNA and lipids of mouse kidney tissues.**

| Raman shift ($cm^{-1}$) | Peak assignment | |
|---|---|---|
| | Nucleic acids | Lipids |
| 1366 | | $CH_3$ symmetric stretch (phospholipids) |
| 1372 | T, A, G (ring breathing modes of the DNA/RNA bases) | $CH_3$ stretching |
| 1437 | | $CH_2$ Deformation<br>Lipids |
| 1445 | | $CH_2$ bending mode of lipids |
| 1456 | Deoxyribose<br>Nucleic acid modes | |
| 1575 | Ring breathing modes in the DNA base<br>Nucleic acid modes indicating the nucleic acid content in tissues | |

**Table S3. Assignments for Raman peaks associated with DNA/RNA, lipids, and proteins of mouse brain tissues.**

| Raman shift (cm-1) | Peak assignment | | |
|---|---|---|---|
| | Nucleic acids | Lipids | Proteins |
| 1040 | | | Phenylalanine |
| 1071, 1098 | | C-C stretch of triacylglycerols | |
| 1130, 1133 | | C-C stretch of cholesterol and cholesterol esters | C-N stretch |
| 1182 | Cytosine, guanine, adenine | C-C stretch of lipid | |
| 1200-1300 | | | Amide III |
| 1303 | | CH3, CH2 twisting and torsion of lipid | |
| 1341, 1343 | A or G of DNA | CH3, CH2 wagging (collagen assignment) | |
| 1375 | T, A, G (ring breathing modes of the DNA/RNA bases) | | |
| 1404 | | | CH deformation |
| 1436, 1438, 1439 | | CH2 deformation, lipid, cholesterol | |
| 1455, 1456 | | CH3, CH2 bending of triacylglycerol | C-H deformation |
| 1480–1575 | | | Amide II |
| 1576, 1578 | Nucleic acid mode, guanine, adenine | | |
| 1585 | | | Phenylalanine, hydroxyproline |
| 1600-1700 | | | Amide I |
| 1618, 1619 | | | Tyr, Trp, Phe |
| 1653 | | C=C stretch of fatty acid | |
| 1674 | | C=C stretch of cholesterol and cholesterol esters | |
| 1729, 1748 | | C=O stretch of triacylglycerol | |

#### Supplementary Figures

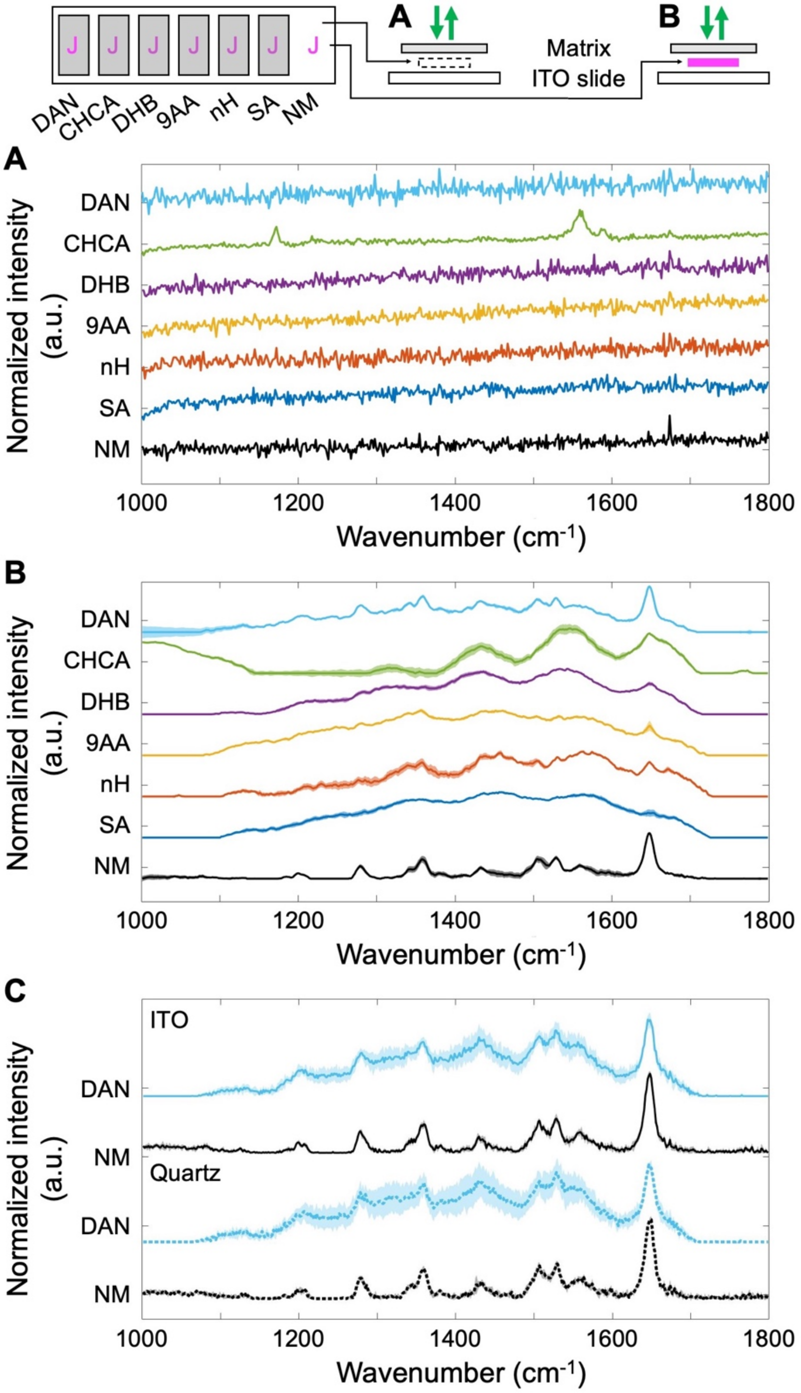

**Figure S1. RaMALDI optimization process to test Raman spectroscopy compatibility with MALDI matrices.** To test and optimize the compatibility of Raman spectroscopy with MALDI matrices, we tested Raman spectra of (**A**) pure MALDI matrices prayed onto ITO slides and (**B**) pink Sharpie® marker drawings on ITO slides sprayed with the six most common MALDI matrices – 1,5-diaminonapthalene (DAN), alpha-cyano-4-hydroxycinnamic (CHCA), 2,5-dihydroxybenzoic acid (DHB), 9-aminoacridine (9AA), norharmane (nH), and sinapic acid (SA) – and without any matrix (NM, no matrix) on ITO slides. (**C**) We then compared spectra of pink Sharpie® marker drawings on ITO slides (top 2 spectra) and quartz slides (bottom 2 spectra), which are standard protocol for Raman measurement both with and without DAN matrix sprayed over the drawings.

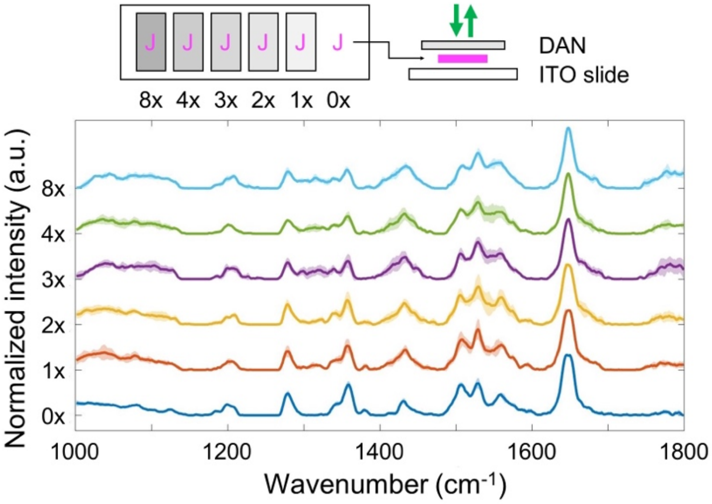

**Figure S2. RaMALDI optimization process to determine matrix density.** To optimize the density of matrix deposition, we tested Raman spectra of pink Sharpie® marker with 1,5-diaminonapthalene (DAN) sprayed at varying density. Number of matrix layers sprayed onto an ITO slide corresponded to matrix density as follows: 0x = 0 mg/mm$^2$, 1x = 0.0004 mg/mm$^2$, 2x = 0.0008 mg/mm$^2$, 3x = 0.0012 mg/mm$^2$, 4x = 0.0016 mg/mm$^2$, and 8x = 0.0032 mg/mm$^2$.

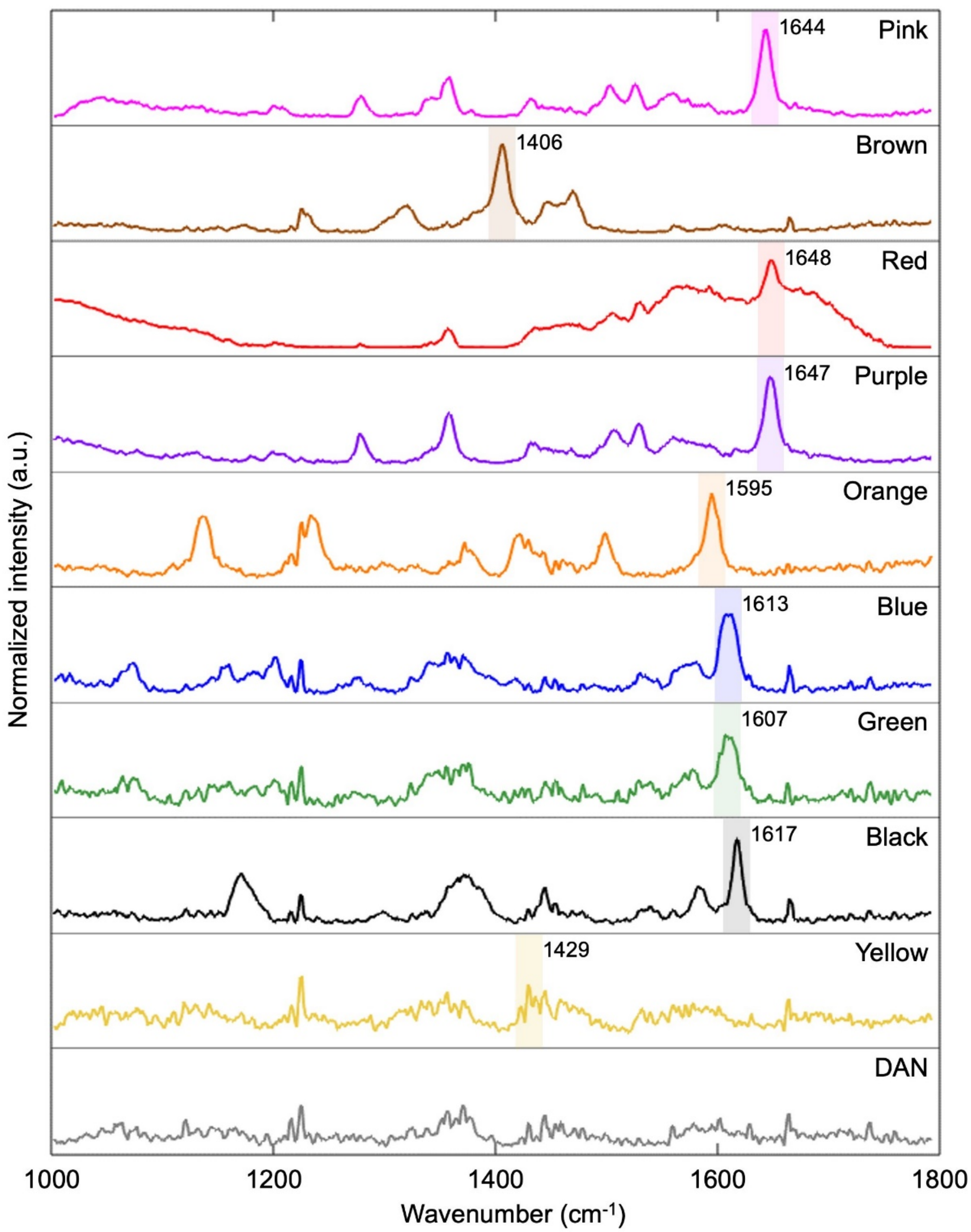

**Figure S3. RaMALDI – Raman spectra of Sharpie® markers of all colors.** Raman spectra of Sharpie® marker drawings on ITO slide sprayed with DAN matrix are shown. Unique Raman peaks (shaded regions) used for evaluating RSI quality and signal scoring as shown in Fig. 2 and Table S1.

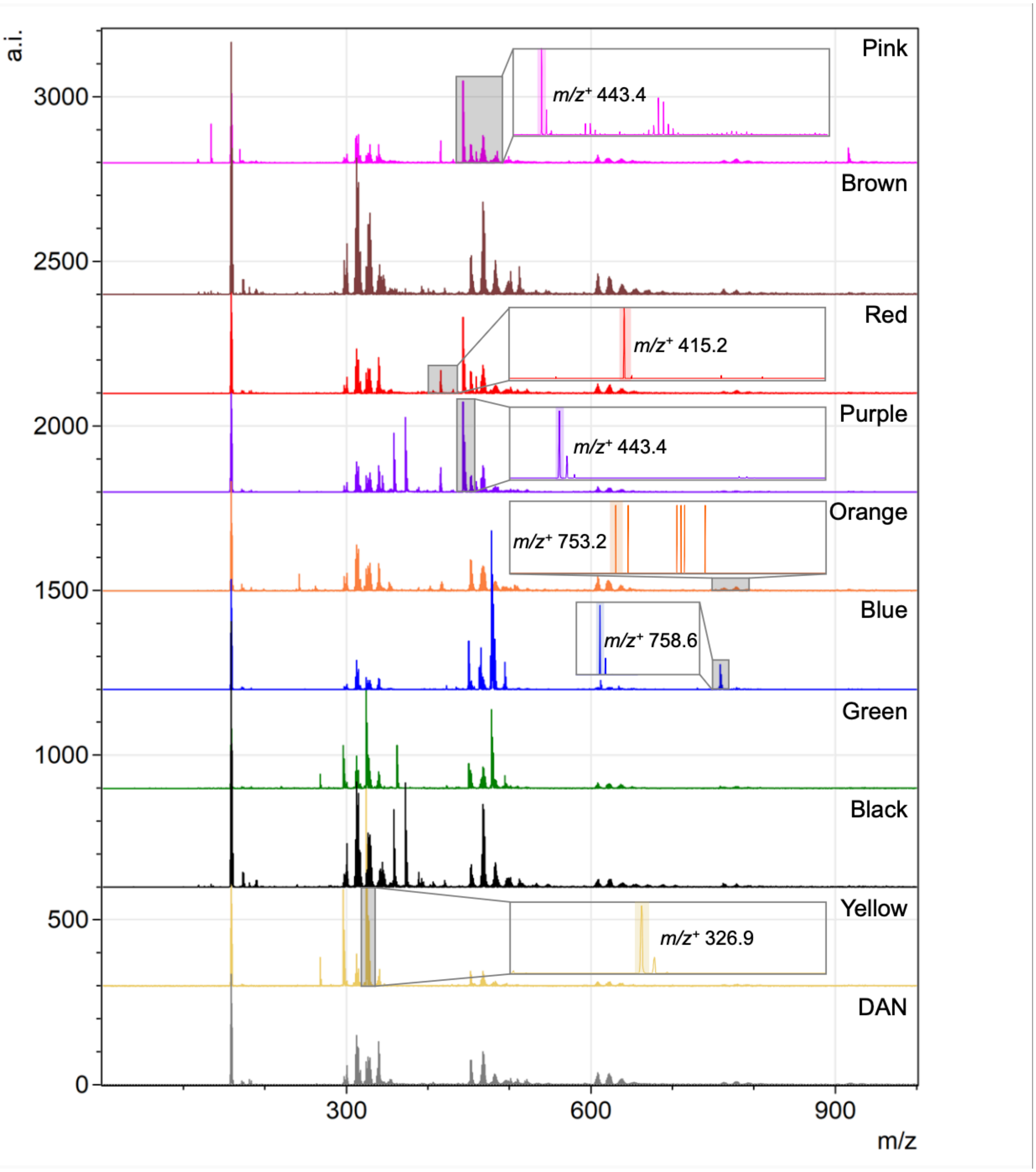

**Figure S4. RaMALDI – positive ion mode MALDI MSI spectra of Sharpie® markers of all colors.** MALDI MSI spectra of Sharpie® marker drawings on ITO slide sprayed with DAN matrix are shown. Unique MS peaks (shaded regions) used for evaluating MALDI MSI quality and signal scoring as shown in Fig. 2 and Table S1.

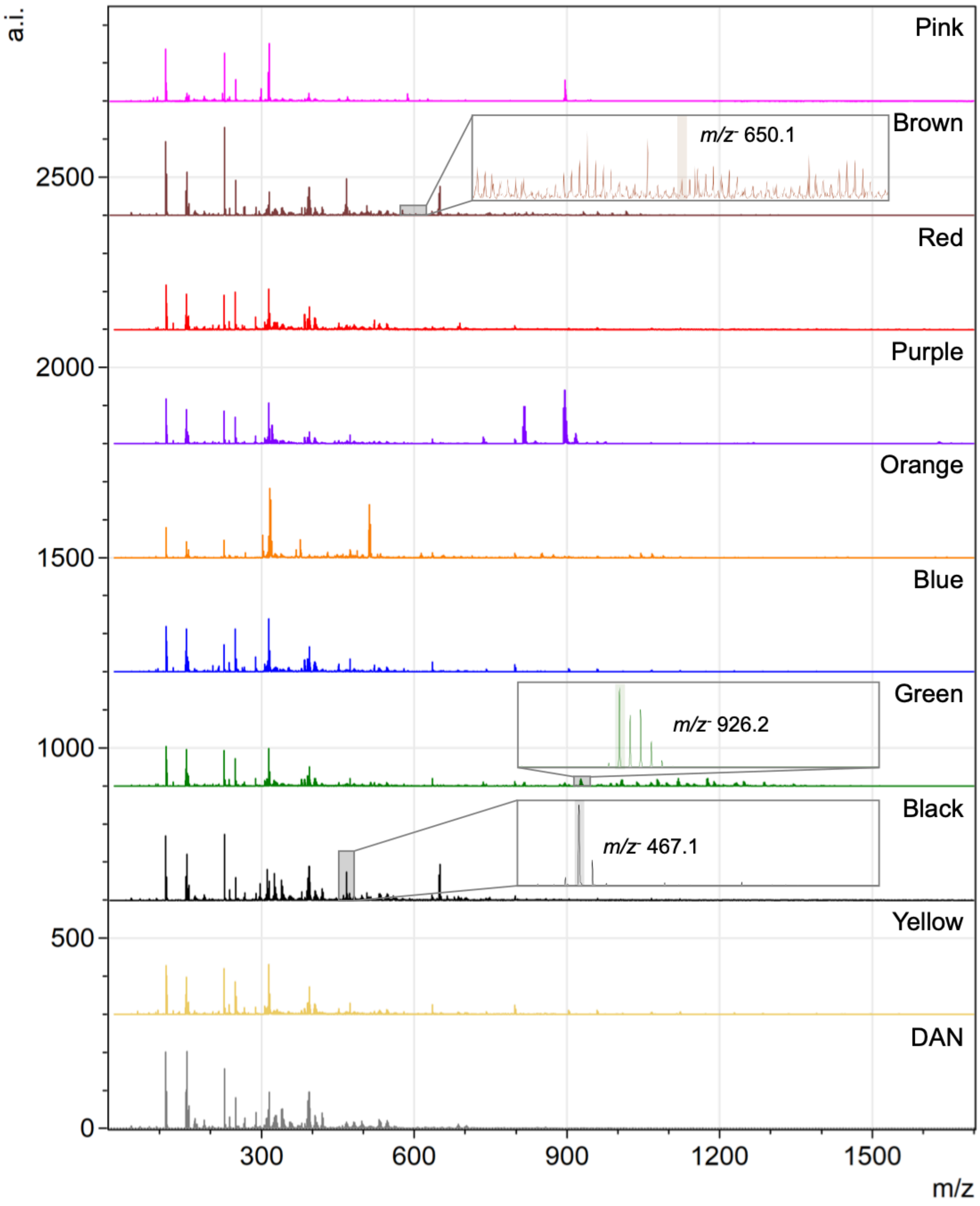

**Figure S5. RaMALDI – negative ion mode MALDI MSI spectra of Sharpie® markers of all colors.** MALDI MSI spectra of Sharpie® marker drawings on ITO slide sprayed with DAN matrix are shown. Unique MS peaks (shaded regions) used for evaluating MALDI MSI quality and signal scoring as shown in Fig. 2 and Table S1.

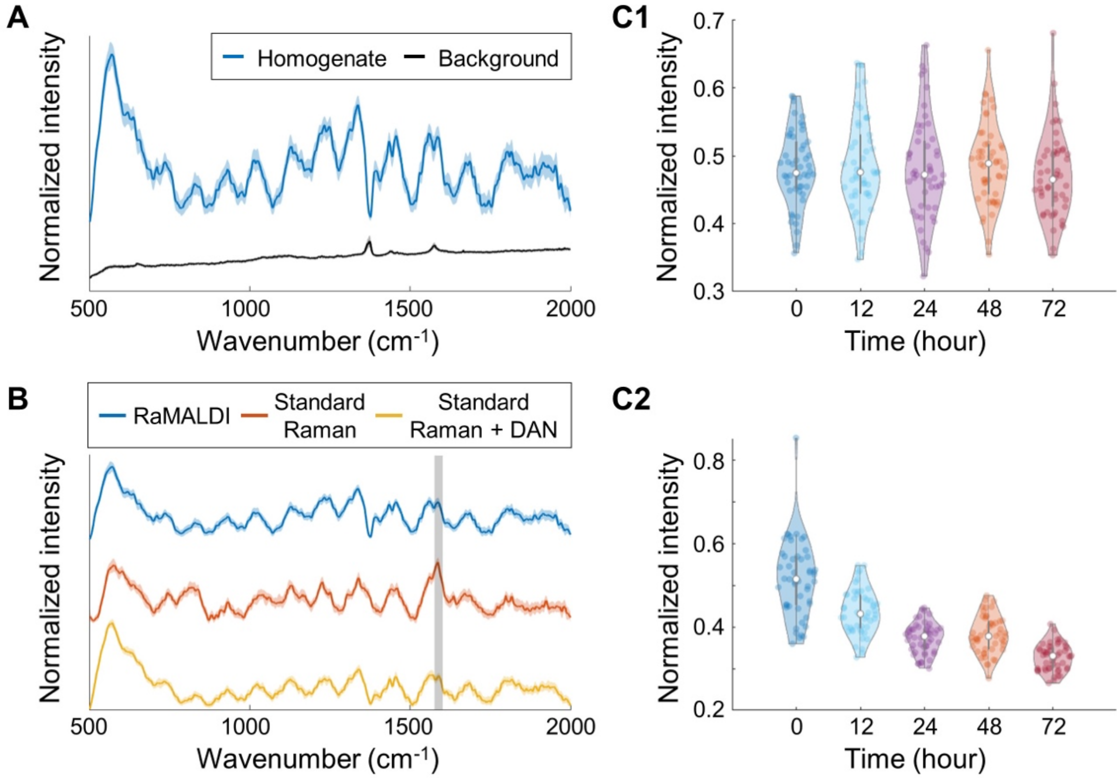

**Figure S6. Sample protection capability of RaMALDI imaging in liver tissue homogenates.** (**A**) RaMALDI - Raman liver homogenate spectra. (**B**) Comparison of Raman liver homogenate spectra measured by RaMALDI imaging (blue), as compared to standard Raman preparation on a quartz slide with (yellow) and without (orange) DAN matrix deposition. (**C**) Changes observed over time in normalized Raman intensity at 1586 $cm^{-1}$ (marked with gray shading in **B**) when preparing the liver homogenate sample with DAN matrix on ITO slide for (**C1**) RaMALDI imaging *versus* without DAN matrix on quartz slide for (**C2**) standard Raman.

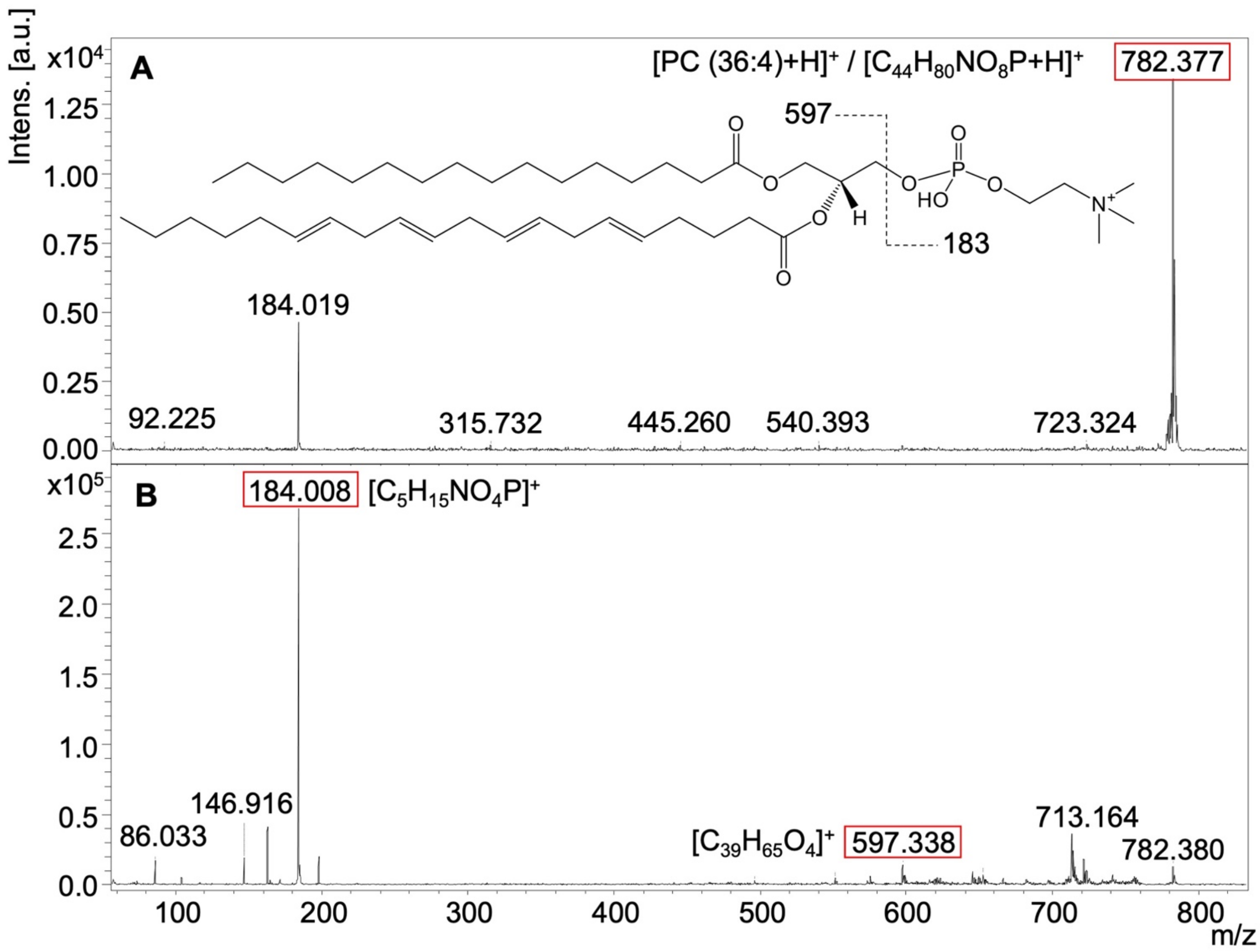

**Figure S7. Positive ion mode MS/MS spectra of m/z⁺ 782.4 identified as PC (36:4), [M+H]⁺.** (**A**) Precursor peak m/z⁺ 782.4, and (**B**) fragmentation of m/z⁺ 782.4. Chemical structures of characteristic fragments (boxed in red) are shown.

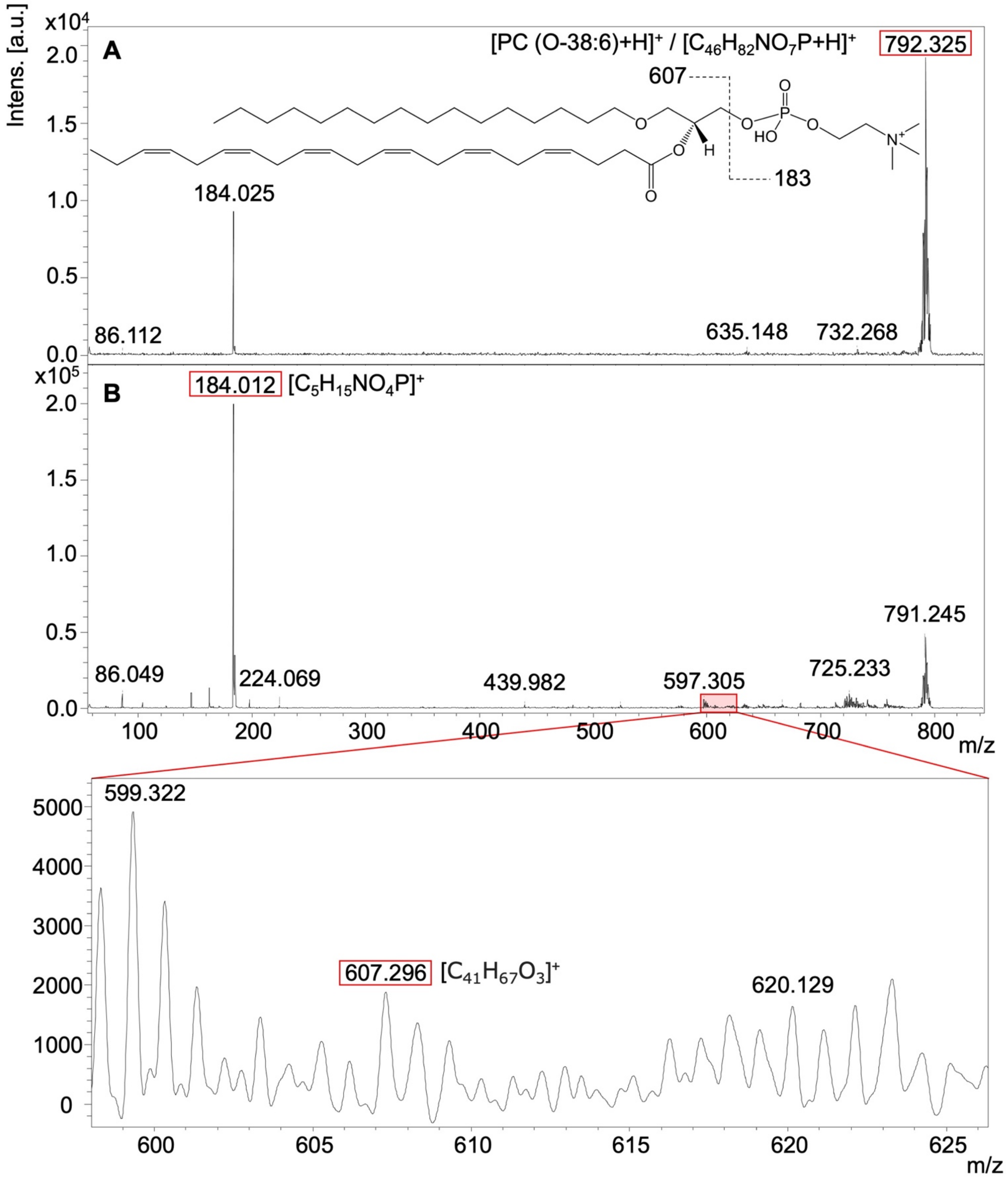

**Figure S8. Positive ion mode MS/MS spectra of m/z$^+$ 792.3 identified as PC (O-38:6), $[M+H]^+$.** (**A**) Precursor peak m/z$^+$ 792.3, and (**B**) fragmentation of m/z$^+$ 792.3. Chemical structures of characteristic fragments (boxed in red) are shown. The phospholipid at m/z$^+$ 792.3 is most likely PC (O-38:6) based on the detected diagnostic ions and high likelihood of even numbered acyl and alkyl chains.

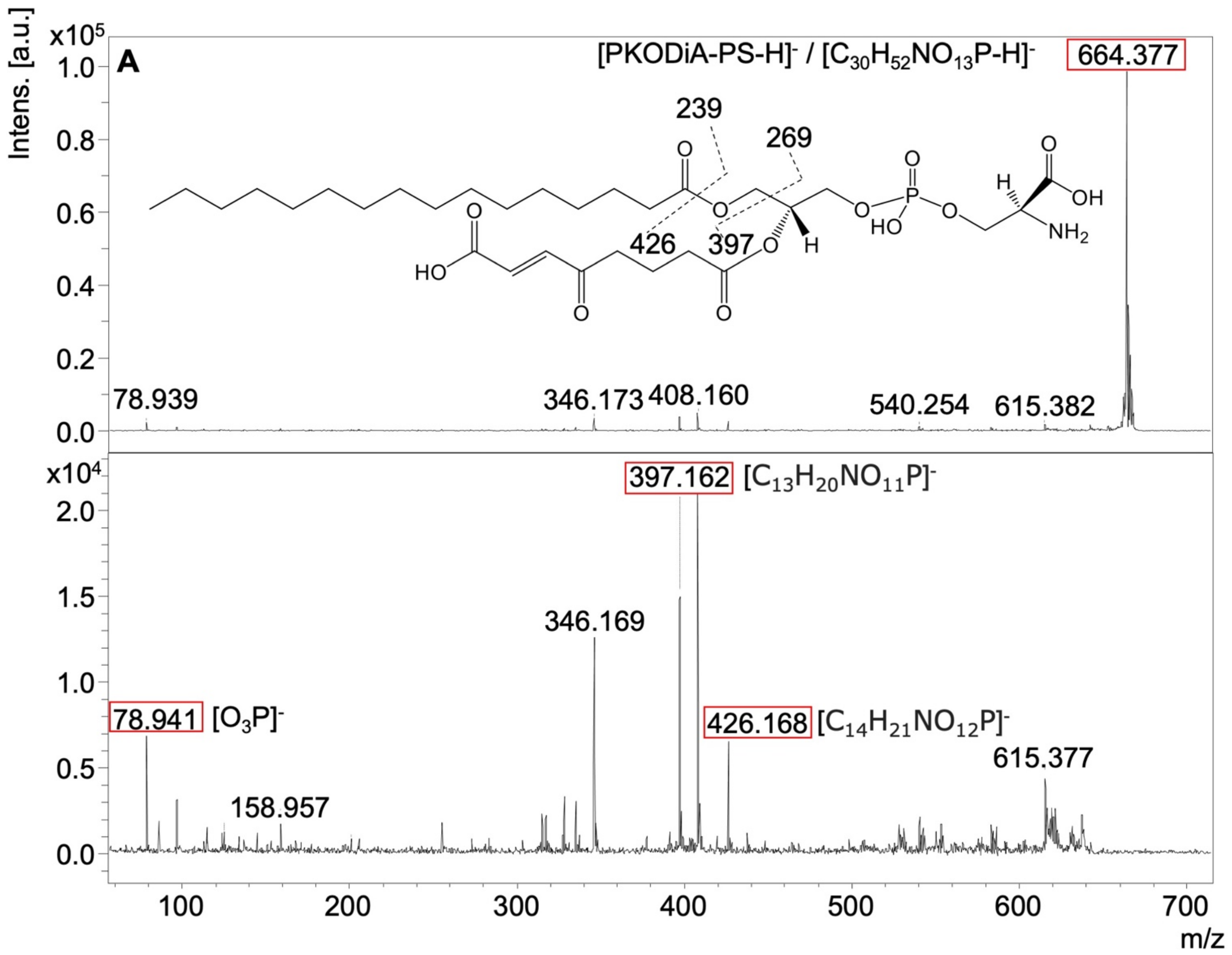

**Figure S9. Negative ion mode MS/MS spectra of m/z⁻ 664.3 identified as PKODiA-PS, [M-H]⁻.** (**A**) Precursor peak m/z⁻ 664.3, and (**B**) fragmentation of m/z⁻ 664.3. Chemical structures of characteristic fragments (boxed in red) are shown.

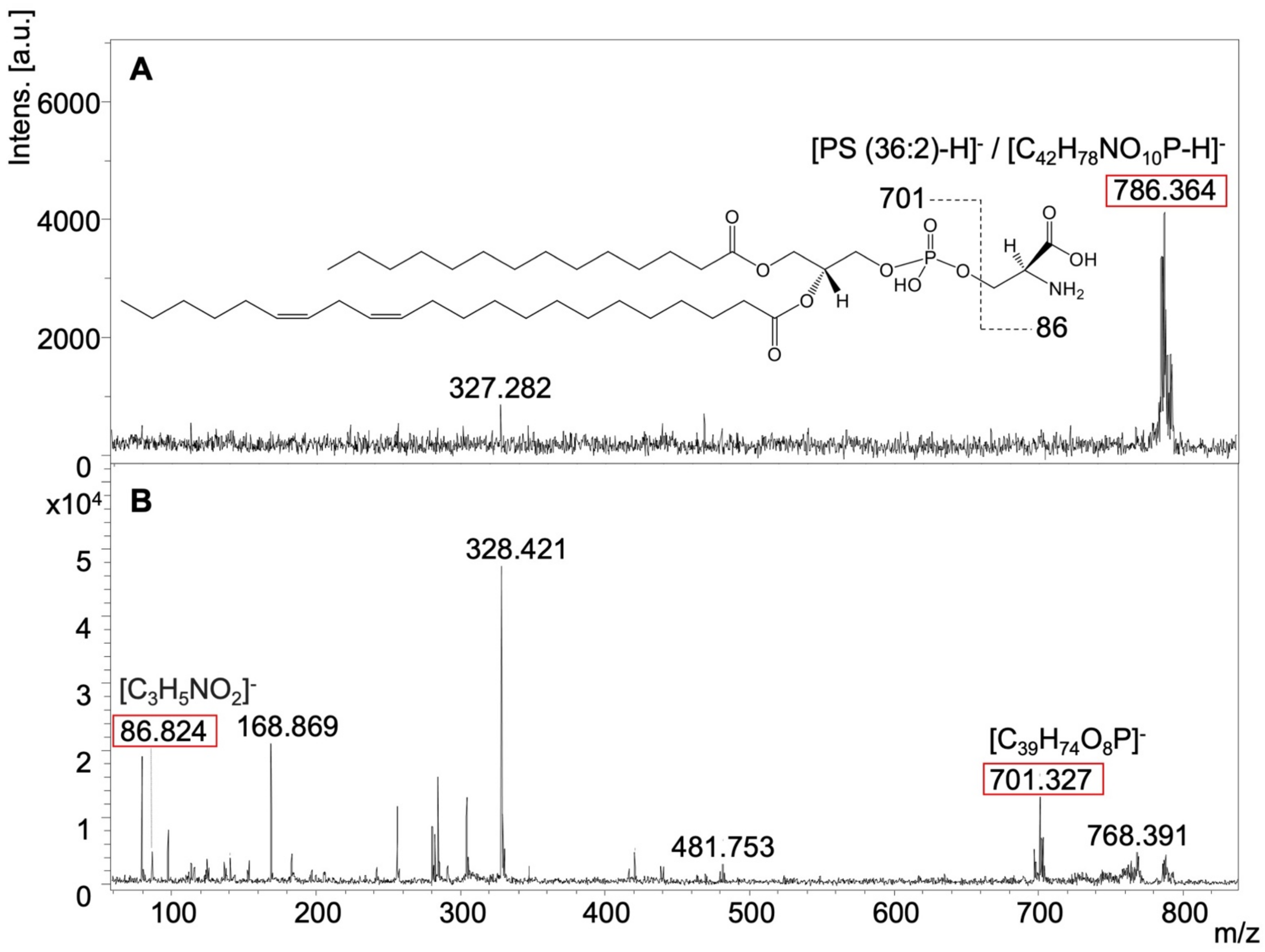

**Figure S10. Negative ion mode MS/MS spectra of m/z$^-$ 786.4 identified as PS (36:2), [M-H]$^-$.** (**A**) Precursor peak m/z$^-$ 786.4, and (**B**) fragmentation of m/z$^-$ 786.4. Chemical structures of characteristic fragments (boxed in red) are shown.

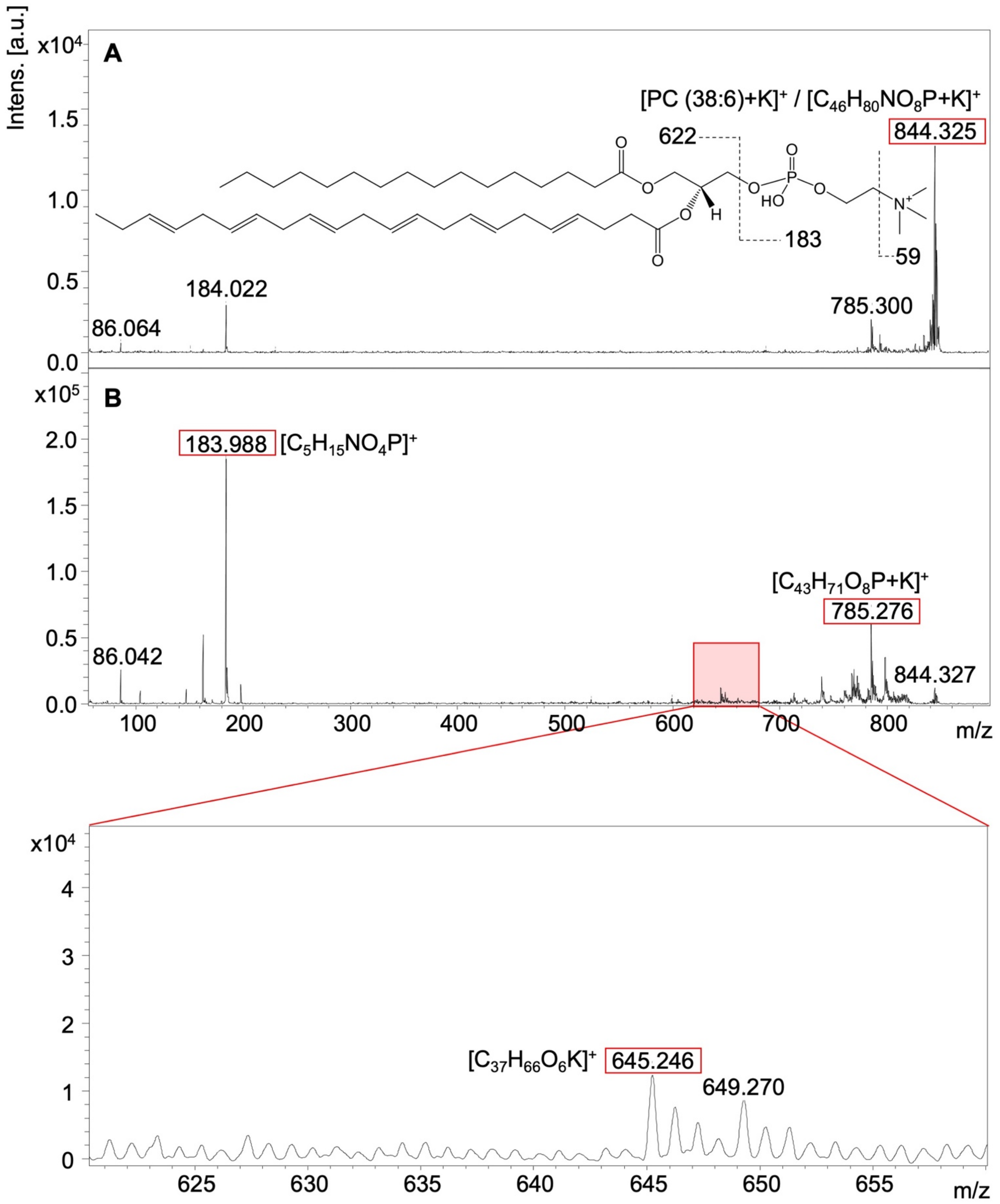

**Figure S11. Positive ion mode MS/MS spectra of m/z+ 844.3 identified as PC (38:6), [M+K]+.** (**A**) Precursor peak m/z+ 844.3, and (**B**) fragmentation of m/z+ 844.3. Chemical structures of characteristic fragments (boxed in red) are shown.

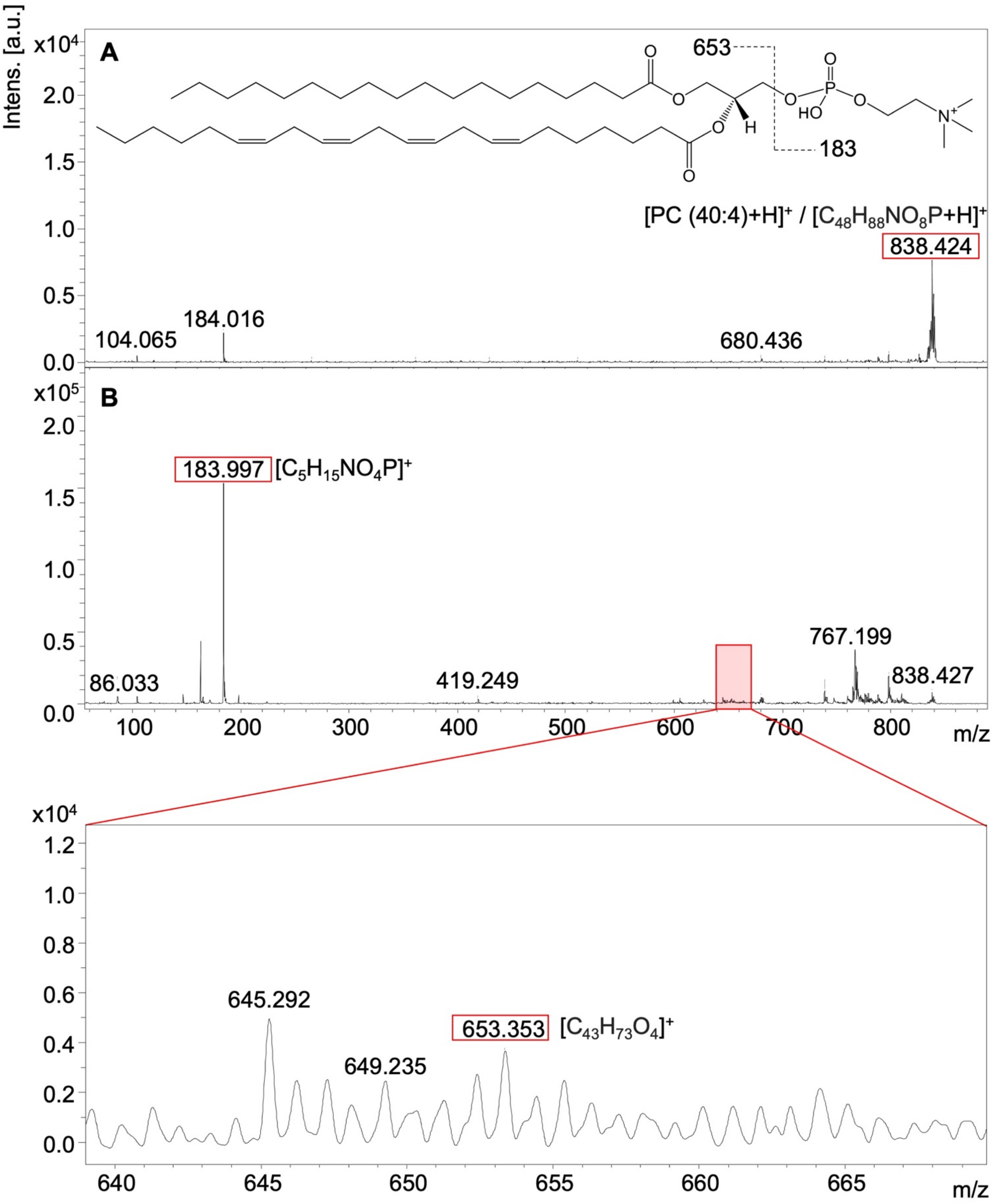

**Figure S12. Positive ion mode MS/MS spectra of m/z⁺ 838.4 identified as PC (40:4), [M+H]⁺.** (**A**) Precursor peak m/z⁺ 838.4, and (**B**) fragmentation of m/z⁺ 838.4. Chemical structures of characteristic fragments (boxed in red) are shown.

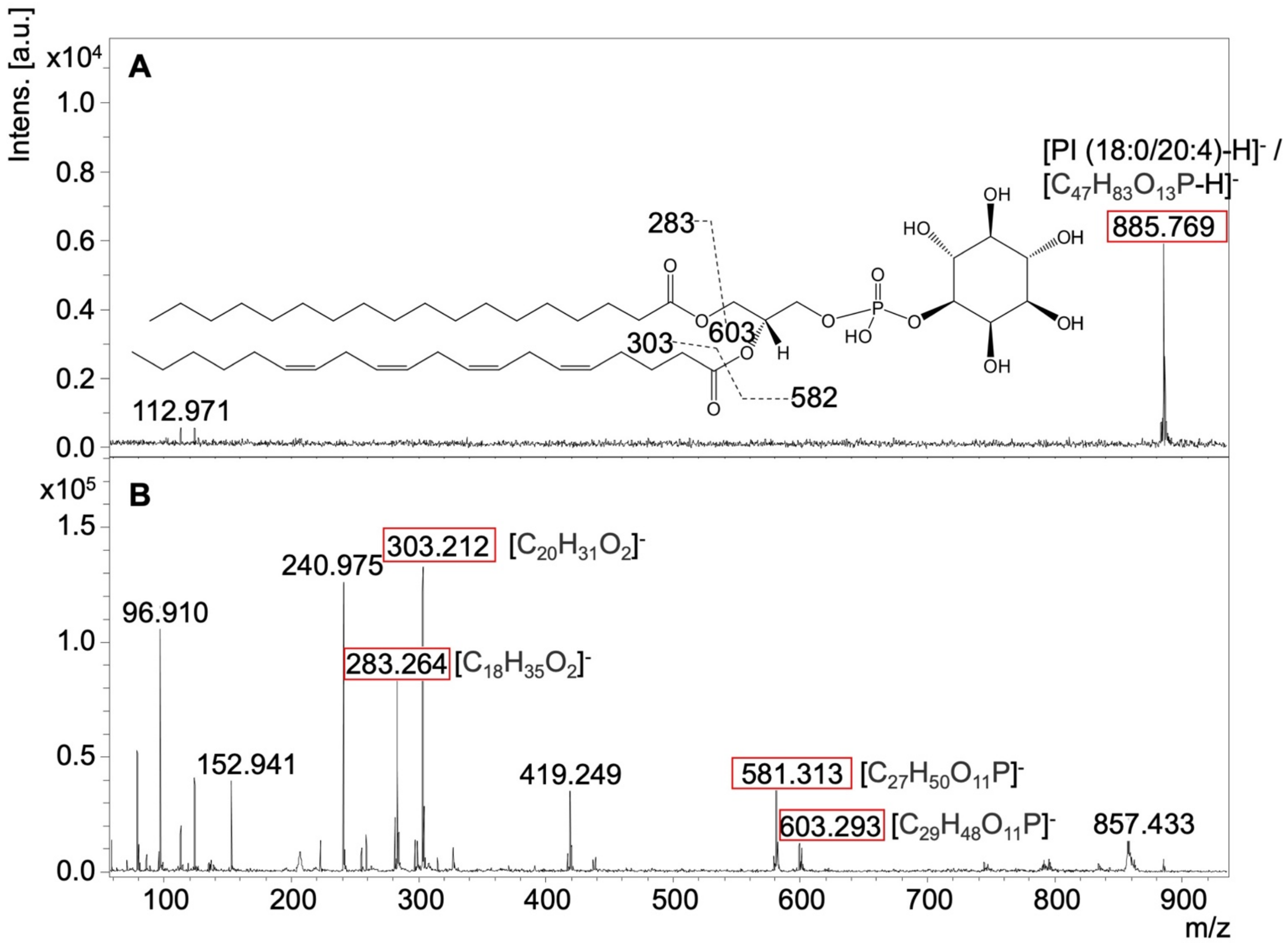

**Figure S13. Negative ion mode MS/MS spectra of m/z⁻ 885.8 identified as PI (18:0/20:4), [M-H]⁻.** (**A**) Precursor peak m/z⁻ 885.8, and (**B**) fragmentation of m/z⁻ 885.8. Chemical structures of characteristic fragments (boxed in red) are shown.

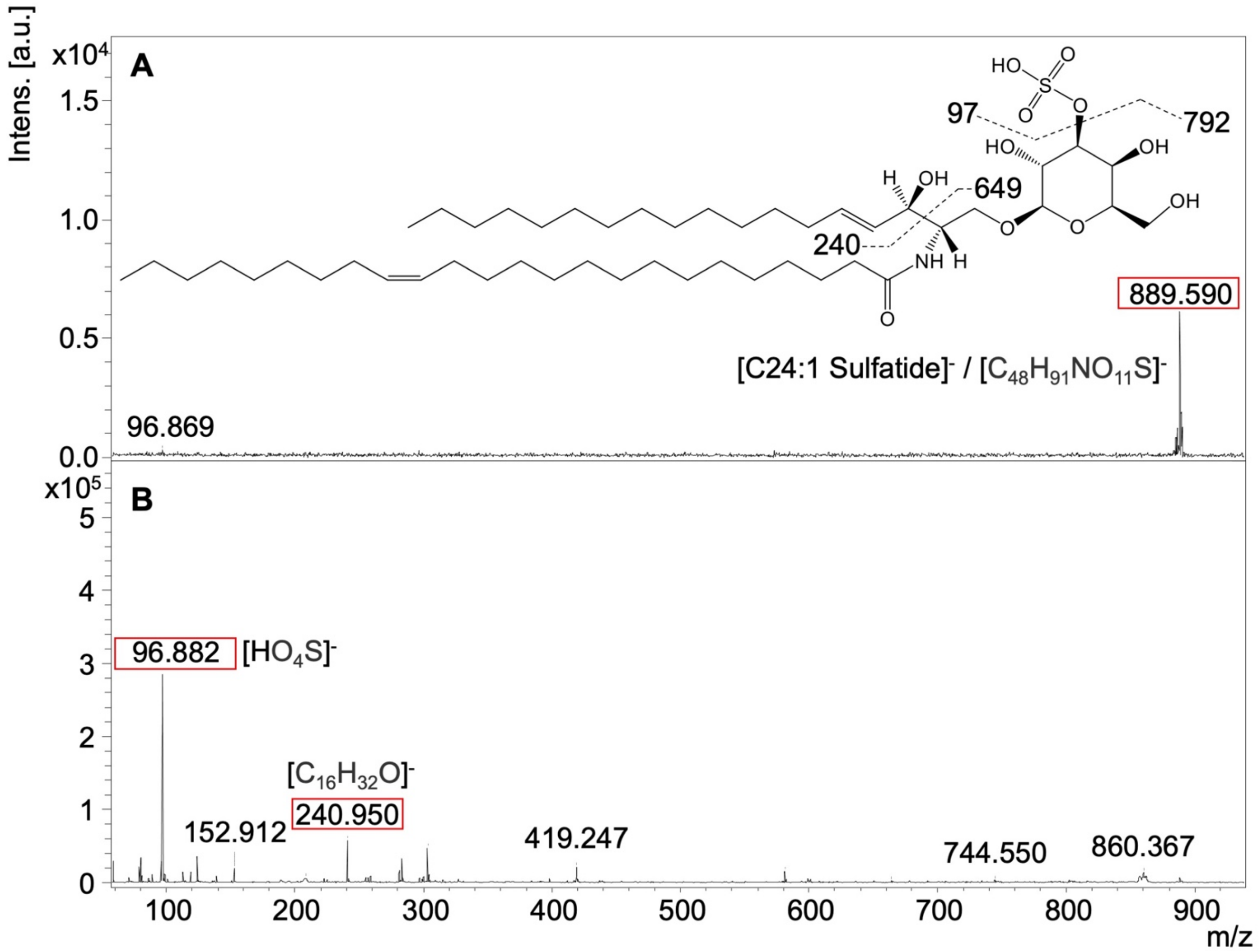

**Figure S14. Negative ion mode MS/MS spectra of m/z⁻ 889.6 identified as C24:1 Sulfatide, [M-H]⁻.** (**A**) Precursor peak m/z⁻ 889.6, and (**B**) fragmentation of m/z⁻ 889.6. Chemical structures of characteristic fragments (boxed in red) are shown.

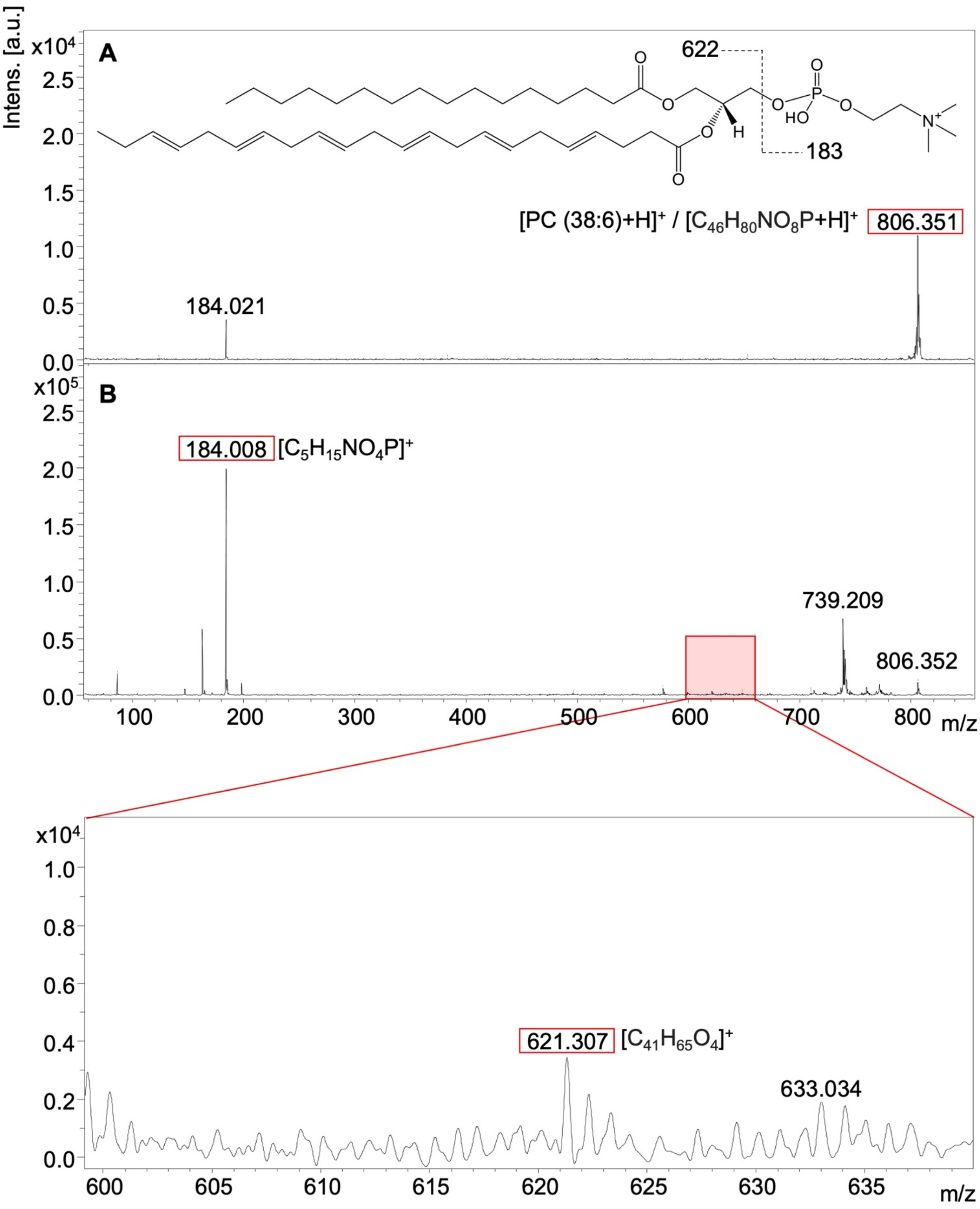

**Figure S15. Positive ion mode MS/MS spectra of m/z$^+$ 806.4 identified as PC (38:6), [M+H]$^+$.** (**A**) Precursor peak m/z$^+$ 806.4, and (**B**) fragmentation of m/z$^+$ 806.4. Chemical structures of characteristic fragments (boxed in red) are shown.

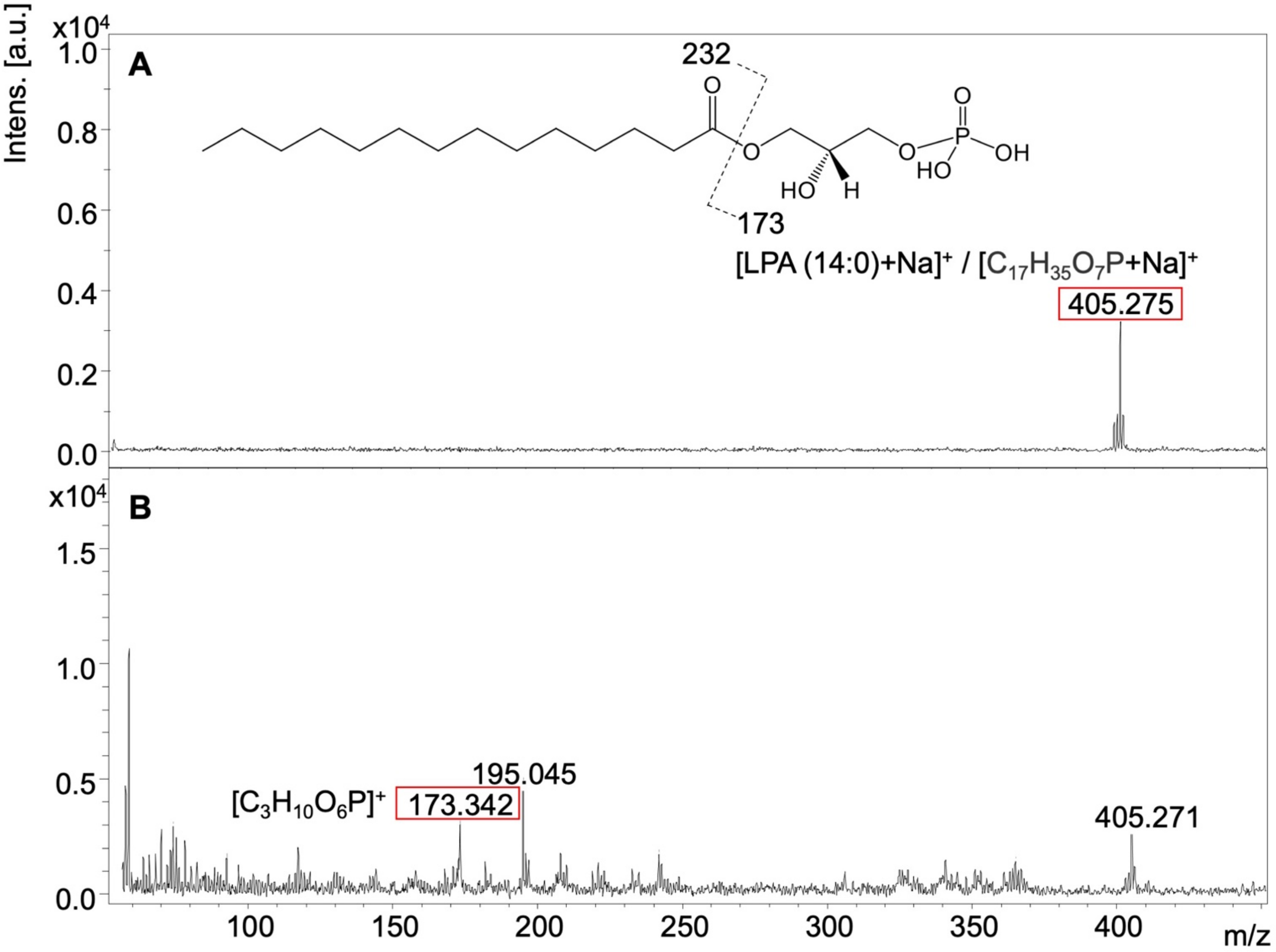

**Figure S16. Positive ion mode MS/MS spectra of m/z⁺ 405.4 identified as LPA (14:0), [M+Na]⁺.** (**A**) Precursor peak m/z⁺ 405.4, and (**B**) fragmentation of m/z⁺ 405.4. Chemical structures of characteristic fragments (boxed in red) are shown.

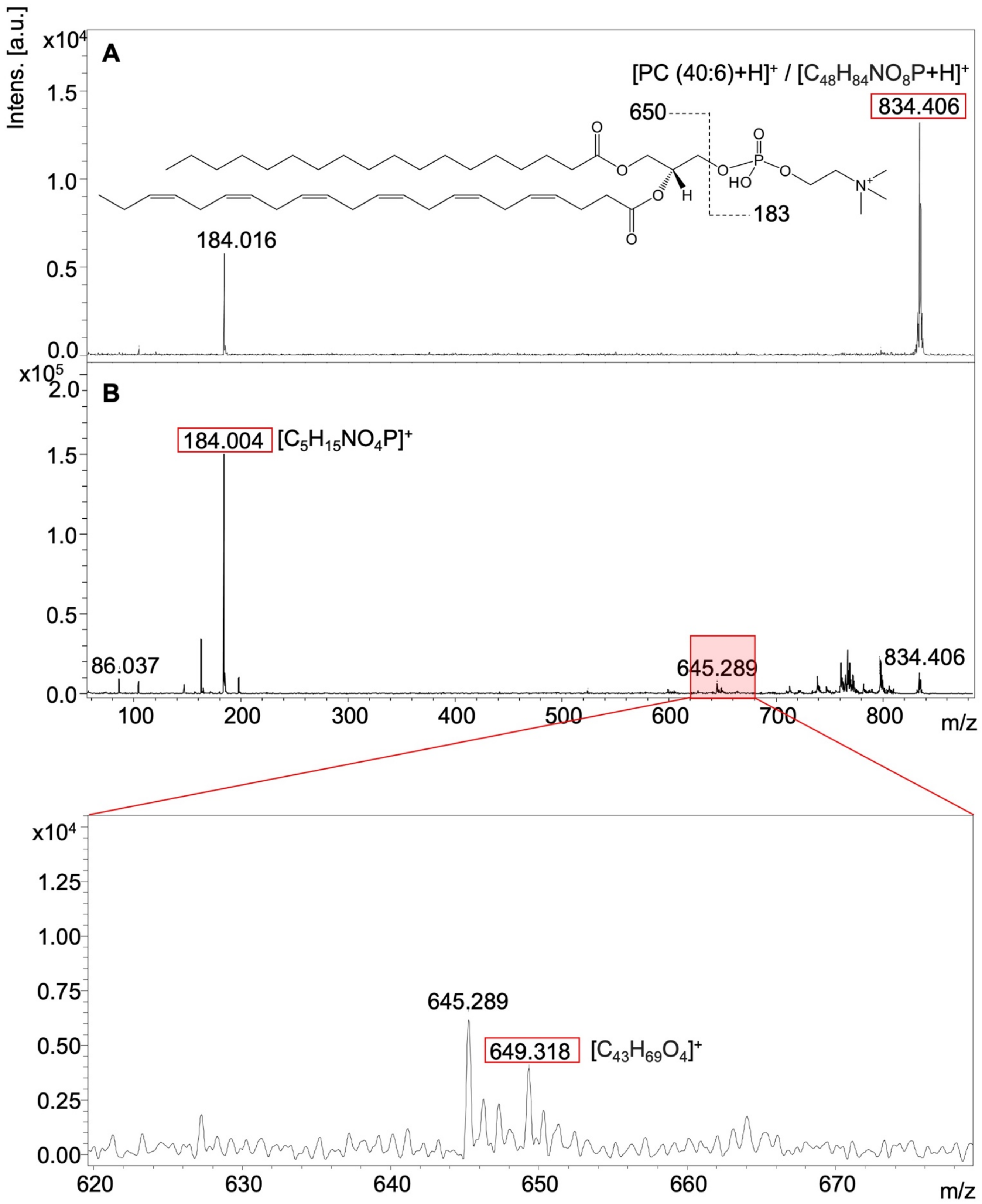

**Figure S17. Positive ion mode MS/MS spectra of m/z+ 834.4 identified as PC (40:6), [M+H]+.** (**A**) Precursor peak of m/z+ 834.4, (**B**) fragmentation of m/z+ 834.4. Chemical structures of characteristic fragments (boxed in red) are shown.

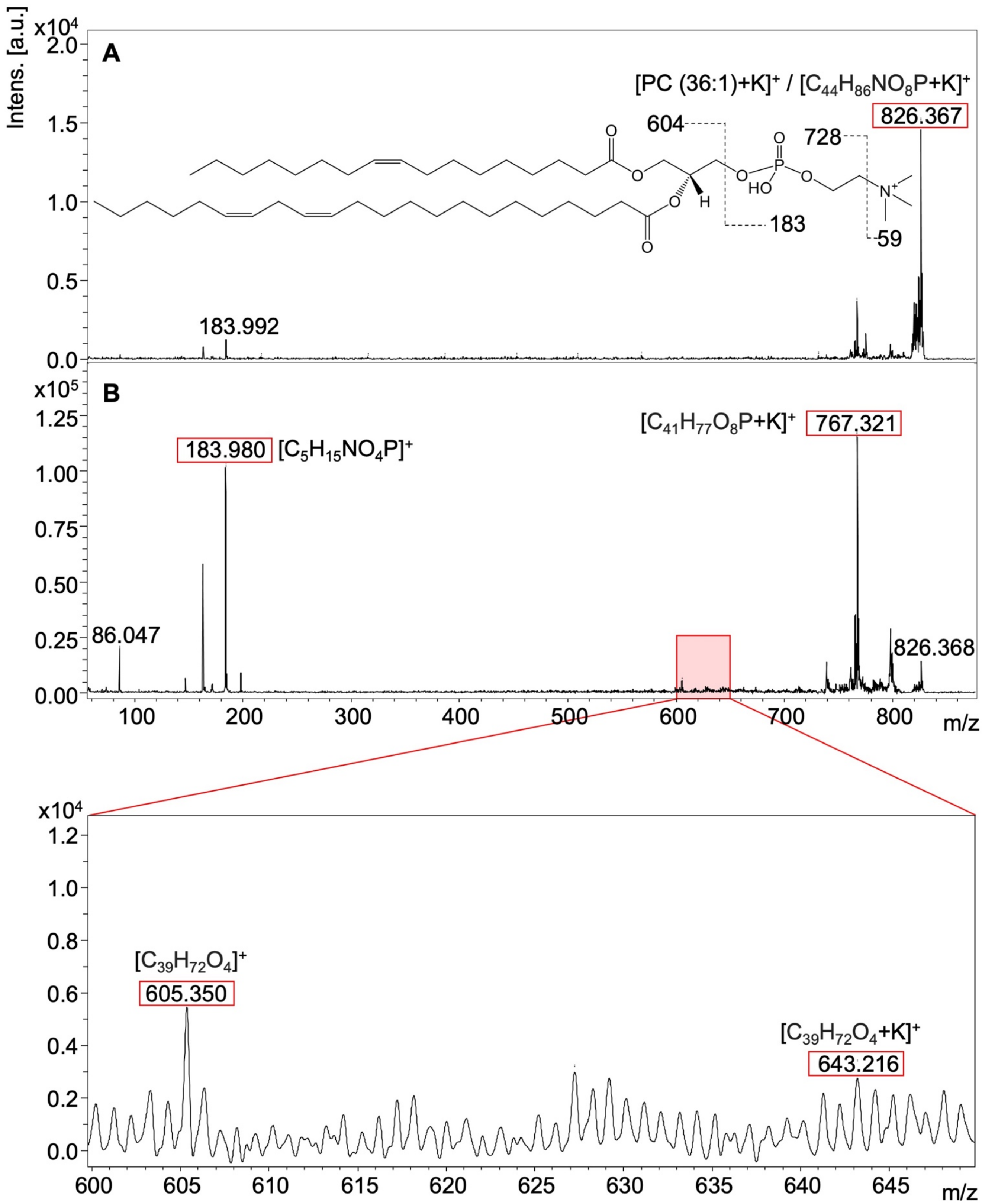

**Figure S18. Positive ion mode MS/MS spectra of m/z+ 826.4 identified as PC (36:1), [M+K]+.** (**A**) Precursor peak of m/z+ 826.4, (**B**) fragmentation of m/z+ 826.4. Chemical structures of characteristic fragments (boxed in red) are shown.

#### Author Contributions

IB and KG conceptualized this study. EY and CMT carried out sample preparation. JHK performed Raman experiments, data processing, and analysis. EY performed MALDI MSI experiments, data processing, and analysis, with assistance from DRB, CCJ, and XES. EY and CMT performed H-E staining. IB and KG directed and supervised the study. JHK, EY, XES, IB, and KG designed the figures. JHK, EY, IB, and KG wrote the manuscript with support from CMT and XES. All authors contributed to, provided critical feedback, reviewed, and approved the final manuscript.
